# Placental insufficiency causes fetal growth restriction in mice lacking *Delta-like homologue 1*

**DOI:** 10.64898/2025.12.16.694688

**Authors:** Maria Lillina Vignola, Ruben Esse, Valeria Scagliotti, Chiara Servadei, Dominika Kardasz, Eugenia Marinelli, Claire Dent, Marika Charalambous

## Abstract

Fetal growth restriction (FGR) affects between 3-7% of pregnancies, is associated with increased perinatal morbidity and mortality, and linked to failure of placental function. The placenta is the key transient organ in pregnancy that directs nutrient transfer, intermediary metabolism and the production of hormones that drive maternal metabolic adaptations essential for pregnancy and lactation. The exchange surface of the placenta is formed in early development by the interaction between trophoblast cells that enclose the maternal blood and extraembryonic mesodermal cells that comprise the fetal vasculature. Despite recent insights into trophoblast development derived from novel in-vitro approaches, the processes driving extraembryonic mesoderm development are not well explored. This is due to a dearth of studies employing unbiased approaches to interrogate extraembryonic mesoderm cell populations. Here we use genetic labelling techniques to separate molecular events occurring in the trophoblast from those in the mesodermal layers of the placenta. In combination with conditional targeting, we show that the imprinted gene *Dlk1* is a key player in providing nutrients to the embryo by controlling the placental surface area available for nutrient exchange, and by modulating the production of placental hormones that promote maternal nutrient provision in pregnancy.

## Introduction

Genomic imprinting describes the process whereby autosomal genes are monoallelically expressed dependent upon their parental origin. The products of imprinted genes are important drivers of fetal growth and placental function (1). Altered dosage of a cluster of imprinted genes on human chromosome 14 causes Temple syndrome, a pediatric growth disorder characterised by intrauterine growth restriction, perinatal faltering growth and alterations to endocrine homeostasis in childhood (2). The chromosome 14 imprinting region contains both paternally-expressed protein coding genes (Delta-like homologue 1 (*DLK1*) and Retrotransposon gag-like 1 (*RTL1*)) and a multi-cistronic maternally-expressed non-coding RNA that includes several functional transcripts (Maternally-expressed gene 3 (*MEG3*), anti-*RTL1*, a large microRNA cluster (*MIRG*) and C/D-box snoRNAs (*RIAN*)). This region is syntenic with distal chromosome 12 in mice (3). Many studies manipulating this imprinted region in mice, including gene-specific deletions, have indicated that the combined actions of genes in the cluster contribute to intrauterine growth restriction and postnatal growth failure (4). However, linking imprinted gene expression to specific molecular pathways of growth and placentation has proved more challenging.

Large genetic screens for human sequence variation associated with birthweight pinpointed a *DLK1*-proximal variant to be associated with low birthweight when inherited from the paternal allele (5, 6). *DLK1* encodes a single-pass transmembrane protein that can be cleaved at the extracellular surface, releasing a 50kDa moiety into the circulation (7). High circulating DLK1 is found in pregnant women, with levels increasing up to term (8). Using *Dlk1*-deleted mice we demonstrated that the source of this maternal circulating DLK1 is the conceptus. In humans, reduced production of DLK1 in the maternal circulation in late gestation is associated with reduced birth weight. Moreover, low circulating DLK1 is linked to small for gestational age (SGA) babies with additional ultrasonic indications of late pregnancy FGR, specifically low umbilical artery pulsatility index at 36 weeks gestational age (wkGA) and reduced abdominal circumference growth velocity between 20-36 wkGA (9). Other groups have since confirmed the link between SGA and low DLK1 (10–12) and linked DLK1 levels to further pregnancy complications such as preeclampsia (13) and maternal insulin resistance (14). Despite this strong relationship between DLK1 levels and intrauterine growth, the molecular underpinnings of these phenotypes are not known.

Since DLK1 is expressed in the placenta of humans and mice, and is associated with FGR in both species, we hypothesised that this protein may have direct actions on placental function. Here, using global and lineage-specific deletion models in mice, we show that *Dlk1* is required in placental endothelial cells for their response to exogenous signalling cues. Loss of *Dlk1* causes failure of endothelial branching morphogenesis and impaired placental growth, highlighting the importance of processes in the extraembryonic mesoderm for correct placental development. Surprisingly, despite the absence of expression in the trophoblast lineage, *Dlk1* deleted placentas altered their production of trophoblast-derived hormones of pregnancy. These hormonal changes were linked to maternal metabolic adaptations associated with adipose storage and energy provision to the fetus. Overall, we conclude that DLK1 is a conceptus-derived regulator of pregnancy, acting directly on placental function to drive both the development of the interhaemal barrier and pathways of fetal-maternal signalling that elicit nutrient exchange.

## Results

### Dlk1 expression is localised to placental cells arising from the extraembryonic mesoderm

To characterise *Dlk1* expression at the single-cell level we used a transcriptomic dataset from isolated single nuclei of the mouse placenta between embryonic days (e)9.5 and e14.5 generated by Marsh et. al. (15). Their initial dimensionality reduction identified four clusters expressing endothelial cell (EC) markers (such as the angiopoietin 1 receptor, *Tek*, and endomucin, *Emcn*) and five that were assigned fetal mesenchyme (FM) identity. *Dlk1* is expressed in three of the four EC clusters (clusters 3, 20 and 25), three of the five FM clusters (clusters 6, 10 and 24), and absent from all other placental cell types (Supplementary Figure 1A-C). Closer examination of markers associated with the original FM clusters indicated that the two *Dlk1*-negative clusters (clusters 15 and 19) may have had their developmental identity misassigned. Based on marker expression, we suggest that these two clusters are likely to represent cells derived from the primitive endoderm lineage. One cluster (cluster 19) expresses *Gata4/6* and parietal endoderm markers such as *Fst*, *Snai1*, *Pth1r*, *Sparc* and *Pdgfra* (*16*), whereas the second (cluster 15) expresses *Gata4/6* in addition to markers of the yolk sac endoderm such as *Afp* and *Ttr* (*17*), (Supplementary Figure 1B). Fetal mesenchymal cells (clusters 6, 10 and 24) all express *Acta2* (encoding smooth muscle actin, SMA). We reassigned cluster 6 as NG2+ pericytes since they express *Cspg4* and *Pdgfrb*, and cluster 24 as stromal cells since they are *Dcn* positive (18). We confirmed the co-localisation of DLK1 with EC and FM markers (EMCN and SMA, respectively, Figure 1A) at e13.5 in the mouse placenta.

**Figure 1.**
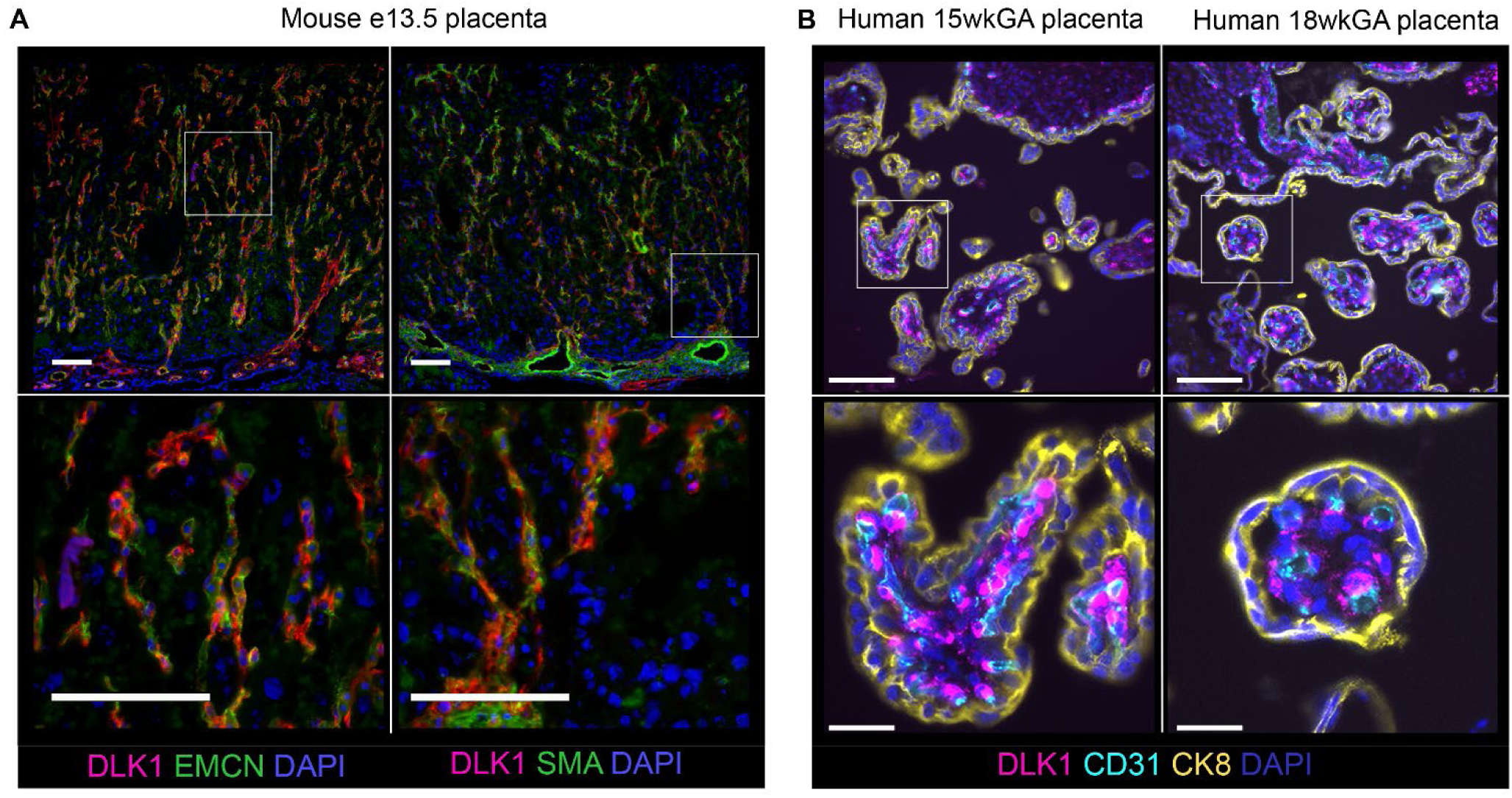
*Dlk1* is expressed in epiblast-derived cells of the mouse and human placenta. A) Immunofluorescence staining of the sagittal-sectioned mouse placental labyrinth at e13.5 with DLK1 (pink) and EMCN (left, green) and SMA (right, green), oriented with maternal decidua at the top and chorionic plate at the bottom. High magnification views (white boxes and images below) indicate cellular co-localisation. All sections are counterstained with DAPI (blue). Scale bar = 100 µm. B) Immunofluorescence staining of human placental villi at 15 (left) and 18 (right) weeks gestational age (wkGA) co-stained with DLK1 (pink) and CD31 (cyan) and cytokeratin 8 (CK8, yellow). All sections are counterstained with DAPI (blue). Scale bar = 100 µm (top) and 25 µm (bottom).

Based on evidence from human single nucleus transcriptomics data (19), *DLK1* localisation appears highly similar between mice and humans. In the second trimester (17-24 wkGA) high levels of expression can be observed in the *ACTA2*/*PDGFRB*-positive mesenchymal cells, and all trophoblast populations are *DLK1*-negative (Supplementary Figure 1D-E). Endothelial cells were poorly represented in these data (Supplementary Figure 1E), so *DLK1* levels could not be robustly ascertained. However, in early (15 wkGA) and mid-second (18 wkGA) trimester human placental villi (Figure 1B), a similar timepoint to the midgestation stage in mice, DLK1 co-localised with the endothelial marker CD31, and also showed abundant expression in the mesenchymal cells within the villi. DLK1 was absent from the syncytial- and cytotrophoblast cells lining the villi (labelled with pan-trophoblast marker cytokeratin 8 (CK8)).

*Dlk1* imprinting was maintained in the mouse placenta over the course of gestation, since transcript levels of *Dlk1* were high when the paternally-inherited allele was intact, but undetectable when the paternally-inherited allele was genetically ablated (Supplementary Figure 2A). Moreover, robust mRNA and protein expression was observed in the labyrinthine zone from mid- to late-gestation placentas with an unmodified paternal copy of the *Dlk1* allele (Supplementary Figure 2B and C, WT and Mat) but not those with a deleted paternal allele (Supplementary Figure 2B and C, Pat and KO).

### Global Dlk1 deletion reduces placental volume by limiting expansion of the labyrinthine zone

To determine the function of *Dlk1* in placental development we generated litters containing individuals with normal *Dlk1* expression or global ablation (Figure 2A) and measured total and compartmental placental volumes between e11.5-e17.5. As expected, placental volume increased in WT animals to a maximum at e15.5. *Dlk1-/-* placentas followed this trajectory up to e13.5, after which placental volume did not increase further (Figure 2B).

**Figure 2.**
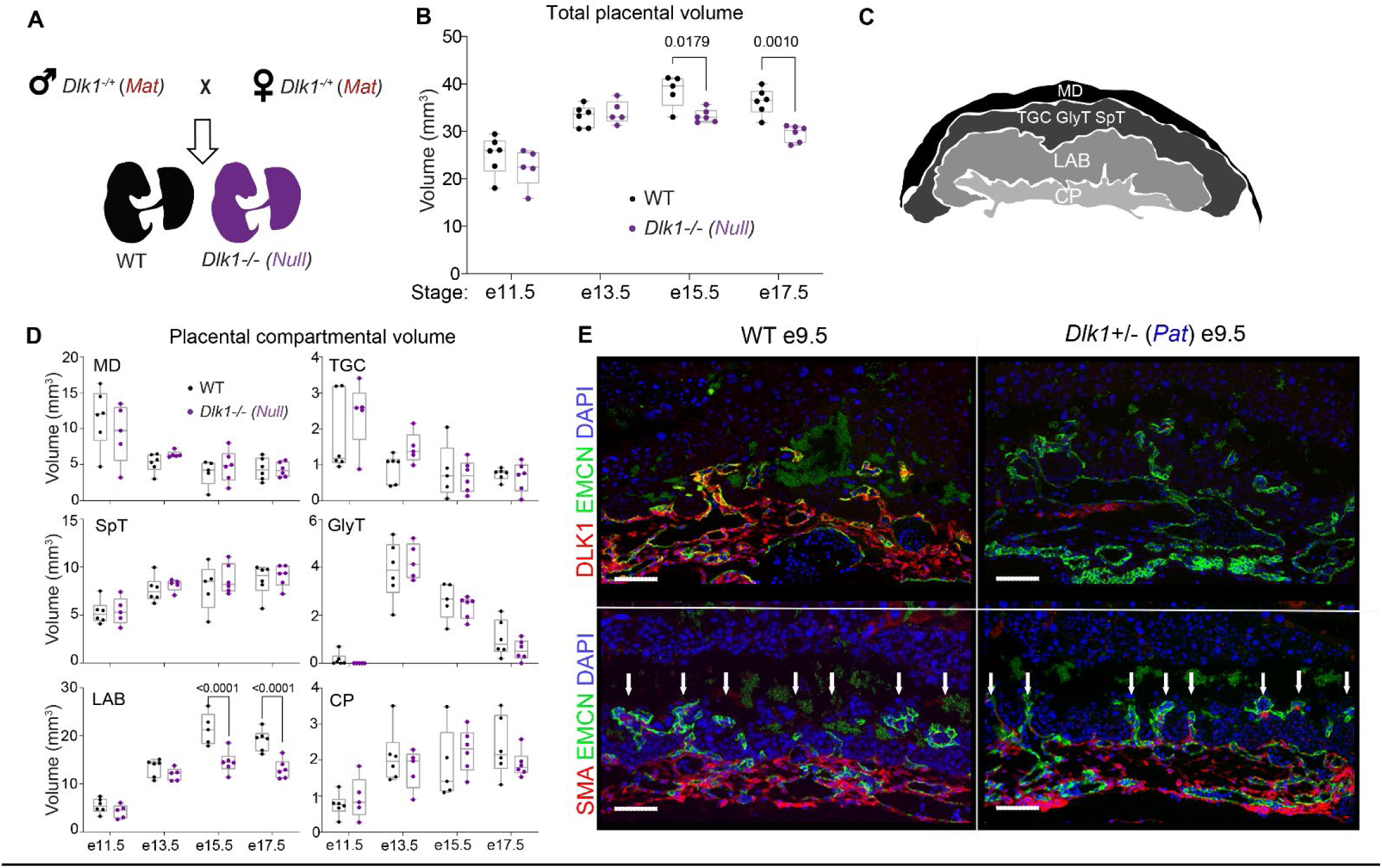
Global loss of *Dlk1* reduces placental volume due to failure of labyrinth expansion in late gestation. A) *Dlk1* maternal heterozygous (Mat) studs and dams were inter-crossed to generate WT and *Dlk1*-Null littermates for analysis of placental volume and compartmental volume by stereology. B) Total placental volume calculated using stereology on WT and *Dlk1*-Null placentas at e11.5 (n = 6 WT, 5 Null), e13.5 (n = 6 WT, 5 Null), e15.5 (n = 5 WT, 6 Null) and e17.5 (n = 6 WT, 6 Null). Data shown in a box and whiskers plot (interquartile range +/− maximum and minimum points in the data). Individual data points, representing individual placentas, are superimposed on the box. Both genotype and stage were significantly different when compared by two-tailed 2-Way ANOVA (p < 0.0001 by stage, p = 0.003 by genotype, p = 0.0202 interaction). Within stages, WT vs Null were compared by Sidak’s multiple comparison test: e11.5 p = 0.453; e13.5 p = 0.987; e15.5 p = 0.0179; e17.5 p = 0.0010. C) Schematic of cell compartments that were quantified using stereology. The top of the image denotes the side of the placenta facing the maternal endometrium, then (top to bottom) maternal decidual cells (MD). The junctional zone contains trophoblast giant cells (TGC), glycogen trophoblast cells (GlyT) and spongiotrophoblasts (SpT). The labyrinth (LAB) zone of the placenta contains highly intercalated cell types. The chorionic plate (CP) marks the embryo-facing side of the placenta. D) Compartmental volumes based on the classification above was estimated across genotypes and stages in samples described in B and compared by two-tailed 2-Way ANOVA. All compartments significantly differed by embryonic stage (MD p < 0.001, TGC p = 0.0002, GlyT p < 0.0001, SpT p < 0.0001, LAB p < 0.0001 and CP p = 0.001). Significant differences between genotypes were seen only in the LAB compartment, and a post hoc Sidak’s multiple comparison test was used to compare WT vs Null placentas at each stage: e11.5 p = 0.863; e13.5 p = 0.615; e15.5 p < 0.0001; e17.5 p = 0.0002). Data is shown in a box and whiskers plots where the box defines the interquartile range and the whiskers show the maximum and minimum points in the data. Individual data points, representing individual placentas, are superimposed on the box, n as in B. E) Immunofluorescence staining of WT (left) and *Dlk1*-Pat (right) sagittal-sectioned mouse placentas at e9.5 co-stained with DLK1 (pink) and EMCN (green, top row). Note a small region of DLK1-positive staining in the *Dlk1*-Pat placenta at this stage which was not observed in subsequent stages. Bottom row: co-staining for EMCN (green) and SMA (pink). White arrows denote EMCN-positive projections from the chorion that form the initial points of endothelial branching. All sections are counterstained with DAPI (blue). Scale bar = 100 µm.

The mature murine placenta is organised into several distinct zones and cell types that can be readily distinguished by haematoxylin and eosin (H&E) staining (20). We classified cells into several types – maternal decidual stroma (MD); trophoblast giant cells (TGC; containing parietal giant cells, spiral artery and channel trophoblast giant cells), glycogen trophoblast cells (GlyT) and spongiotrophoblast cells (SpT) of the junctional zone; labyrinthine zone cells (LAB, comprised of closely interdigitated trophoblast cells with fetal endothelial and mesenchymal cells), and chorionic plate cells (CP) (Figure 2C). By combining stereological analysis with cell type identification we found that the reduction in placental volume in *Dlk1-/-*placentas was primarily due to a reduction in the size of the labyrinthine layer (Figure 2D, LAB), which was ∼50% reduced in size in the late gestation placenta in the absence of *Dlk1*.

The labyrinth of the placenta constitutes the interface by which nutrients and gases are exchanged between the maternal and fetal circulation. The formation of this interface initiates between e9-e10 of gestation, when the trophoblast-derived cells of the chorion are contacted by the allantois, an extraembryonic structure derived from the mesoderm. Endothelial cells of the allantois form projections into the chorionic plate and thereafter expand and undergo branching morphogenesis. In concert, the trophoblast expands and differentiates into specialised cells that will encapsulate the maternal blood. By e13.5 a highly interdigitated trilaminar structure has formed that separates the maternal and fetal blood cells (21). DLK1 is expressed at high levels in the extraembryonic mesoderm at e9.5, including in EMCN-positive endothelial cells projecting into the chorionic plate (Figure 2E). Loss of *Dlk1* expression does not impair chorioallantoic attachment since EMCN-positive endothelial projections infiltrate the chorionic trophoblast layer in *Dlk1*-mutant placentas with similar frequency to the wild-type (Figure 2E). Moreover, both WT and *Dlk1*-mutant placentas have clearly differentiated endothelial and SMA-positive mesenchymal cell populations, indicating that the early stages of chorioallantoic attachment and vasculogenesis are not impaired by loss of *Dlk1*.

### Global Dlk1 deletion impairs endothelial development

From the initiation of chorioallantoic attachment to the end of gestation, the labyrinthine zone of the mouse placenta becomes progressively more complex. Fetal endothelial cells derived from the extraembryonic mesoderm are first observed in wide projections that intercalate with large islands of labyrinthine trophoblast cells (e9.5-e10.5). This can be visualised by co-staining with EMCN and the pan-trophoblast marker cytokeratin 7 (CK7, Figure 3A-B). By e11.5, the EMCN-positive cells have formed clearly branched vessels that contain nucleated fetal blood cells, and by e13.5 the labyrinth has formed a dense structure with highly interdigitated endothelium and trilaminar trophoblast that retains the maternal circulation (Figure 3C-D). Alongside the progression of branching morphogenesis of the fetal endothelial cells, SMA-positive mesenchymal cells leave the chorionic plate region and colonise the labyrinthine zone, forming tight contacts with the endothelial cells (Figure 3E-G).

**Figure 3.**
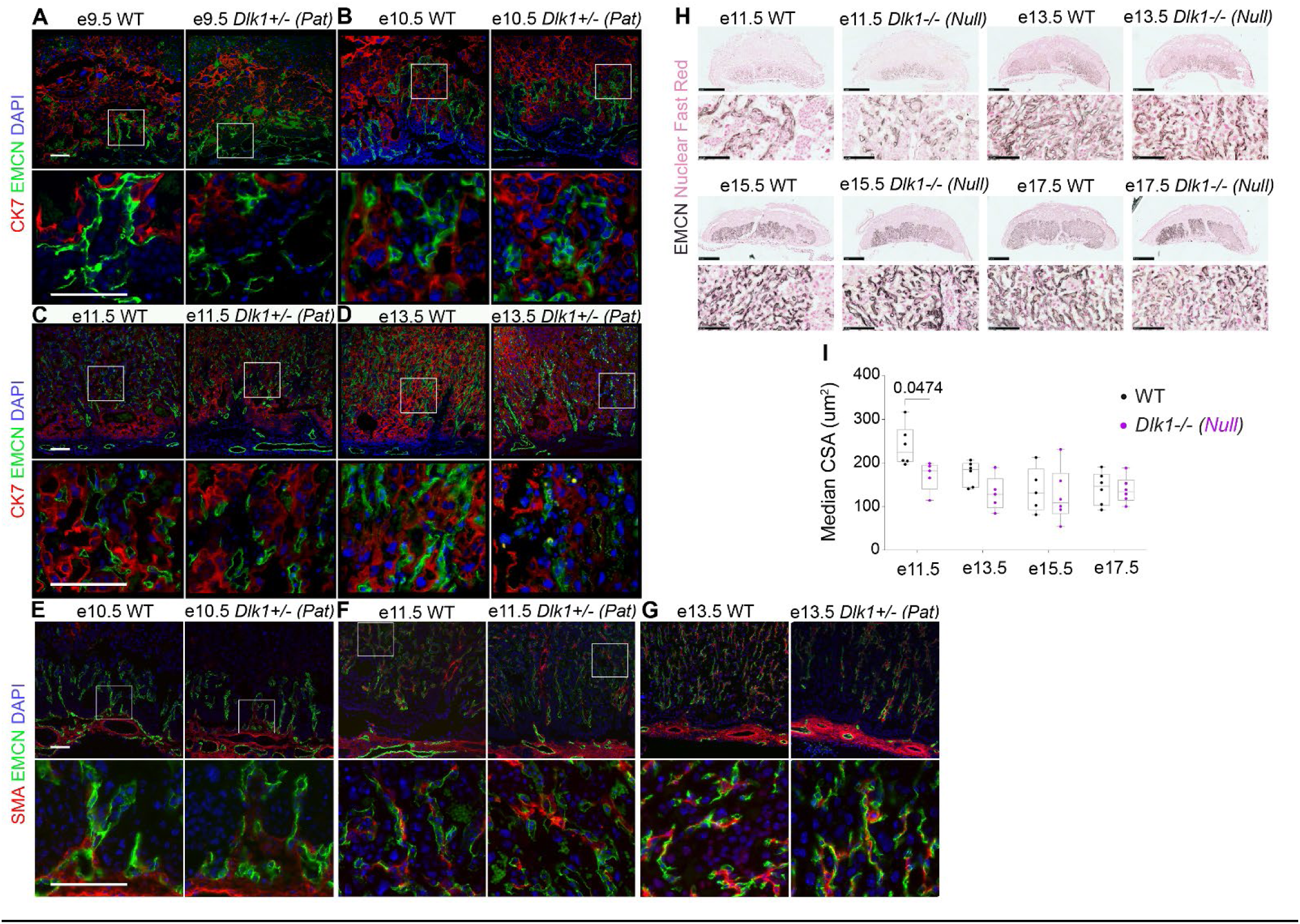
Loss of Dlk1 impairs formation of the placental labyrinthine endothelium. A-D) Immunofluorescence staining of WT (left) and *Dlk1*-Pat (right) sagittal-sectioned mouse placentas at e9.5 (A), e10.5 (B), e11.5 (C), e13.5 (D) co-stained with CK7 (pink) and EMCN (green). Images below show a high magnification view of the boxed section. For all images the scale bar = 100um. Images are counterstained with DAPI (blue). E-G) Immunofluorescence staining of WT (left) and *Dlk1*-Pat (right) sagittal-sectioned mouse placentas at e10.5 (E), e11.5 (F), e13.5 (G) co-stained with SMA (pink) and EMCN (green). Images below show a high magnification view of the boxed section (except for G where the boxed section lies outside the larger image). For all images the scale bar = 100 µm. Images are counterstained with DAPI (blue). H) Immunohistochemistry for EMCN at embryonic stages e11.5, e13.5, e15.5 and e17.5 in sagittally-sectioned mouse placentas derived from heterozygous inter-crosses of *Dlk1*-deleted parents, WT and *Dlk1*-Null littermates. Bottom panels show the labyrinth region of the placenta. Positive staining is stained in black and counterstained with nuclear fast red (pink). Scale bar shows 1 mm in the top panels and 100 µm in the bottom panels. I) Endothelial cross sectional area (CSA) was estimated from EMCN staining as in H, on WT and *Dlk1*-Null placentas from Figure 2, at e11.5 (n = 6 WT, 5 Null), e13.5 (n = 6 WT, 5 Null), e15.5 (n = 5 WT, 6 Null) and e17.5 (n = 6 WT, 6 Null). Data is shown in a box and whiskers plot where the box defines the interquartile range and the whiskers show the maximum and minimum points in the data. Data points, representing median CSA values from individual placentas, are superimposed on the box. Both genotype and stage were significantly different when compared by two-tailed 2-Way ANOVA (p = 0.0012 by stage, p = 0.0145 by genotype). Within stages, WT vs Null were compared by Sidak’s multiple comparison test: e11.5 p = 0.0474; e13.5 p = 0.283; e15.5 p = 0.983; e17.5 p = 0.999.

Placentas lacking *Dlk1* showed impairments in this process that precede the changes in total and labyrinth volume (Figure 2B, D). From e9.5-e10.5 WT and *Dlk1*-Pat placentas showed a similar pattern of endothelial infiltration into the trophoblast bed (Figure 3A-B). By e11.5, *Dlk1*-Pat placentas had noticeably fewer EMCN-positive vessels, and by e13.5 the mutant placenta had much sparser distribution of endothelial vessels and expanded areas that were solely populated by trophoblast (Figure 3C, D). Similarly, the distribution and abundance of mesenchymal cells did not appear different between WT and *Dlk1*-Pat placentas up to e11.5 (Figure 3E, F), but by e13.5 SMA-positive mesenchymal cells in proximity to the labyrinthine endothelium appeared reduced in abundance (Figure 3G).

As endothelial branching progresses, vessel cross sectional area (CSA) is reduced (20). We observed a marked decrease in EMCN-positive vessel CSA between e11.5 and e15.5 in WT placentas (Figure 3H, I), coinciding with the known increase in branching morphogenesis in this interval (20). However, in placentas lacking *Dlk1* endothelial CSA was already reduced at e11.5 and appeared to reach the steady-state mature vessel diameter earlier than the WT (e13.5 in mutants vs e15.5 in WT, Figure 3H, I). Taken together these data suggest that in the absence of functional *Dlk1* the morphogenesis of the placental labyrinth is impaired such that vascular structures mature prematurely and by late gestation fail to intercalate with the trophoblast at high density.

### Expression of Dlk1 in endothelial cells is required for placental and fetal growth

We and others have previously demonstrated that global deletion of *Dlk1* in mice causes placental and embryonic growth restriction in late gestation (9, 22, 23). Since *Dlk1* is expressed at high levels in the endothelial population of the placenta (Figure 1) and the labyrinth appeared most affected by its deletion, we asked if deleting *Dlk1* in the endothelial compartment is sufficient to recapitulate the *Dlk1* global deletion phenotype. Three lines of *Dlk1*-deleted mice were generated, then embryonic and placental mass was compared at e15.5 (Figure 4A). The global deletion of *Dlk1* at this stage showed the expected ∼10% reduction in fetal and placental mass (Figure 4B, C). Male and female *Dlk1*-deleted embryos exhibited a similar level of growth restriction of both embryo and placenta. In addition, female placentas were smaller than male placentas (Supporting Figure 3A, B). Next, we generated conceptuses that lack *Dlk1* in all cells derived from the epiblast (Figure 4A), using a conditional *Dlk1* (floxed) allele (23) and a *Meox2*-Cre driver (24). Since *Dlk1* is not expressed in the WT trophoblast, this model should recapitulate the global deletion. This was indeed the case, since the epiblast-specific *Dlk1* deleted (*Dlk1*-EB) conceptuses also had ∼10% reduction in fetal and placental mass (Figure 4B, C). Finally, we used a *Tek*-Cre driver (25) to determine the influence of endothelial cell expression of *Dlk1* on placental phenotype (25). We confirmed that this model (*Dlk1*-VE, Figure 4A) had deleted *Dlk1* in the placental endothelial cells, since DLK1 was no longer co-localised with EMCN in the *Dlk1*-VE placenta (Supplementary Figure 4A). However, we observed an unexpected increase in DLK1 expression in the unmanipulated mesenchymal population, which was confirmed by real-time quantitative PCR (Supplementary Figure 4B, C). When *Dlk1* was deleted from the vascular endothelium using this line (*Dlk1*-VE), we observed a ∼7% reduction in embryonic mass and a ∼18% decrease in the mass of the placenta (Figure 4B, C). As with the global *Dlk1* deletion, *Dlk1*-VE placentas displayed impaired morphology of the placental vasculature, including reduced endothelial density and mesenchymal cell abundance (Figure 4D, E). These data are consistent with a cell-autonomous role for *Dlk1* in the placental vasculature to regulate placental labyrinth morphology and fetal growth.

**Figure 4.**
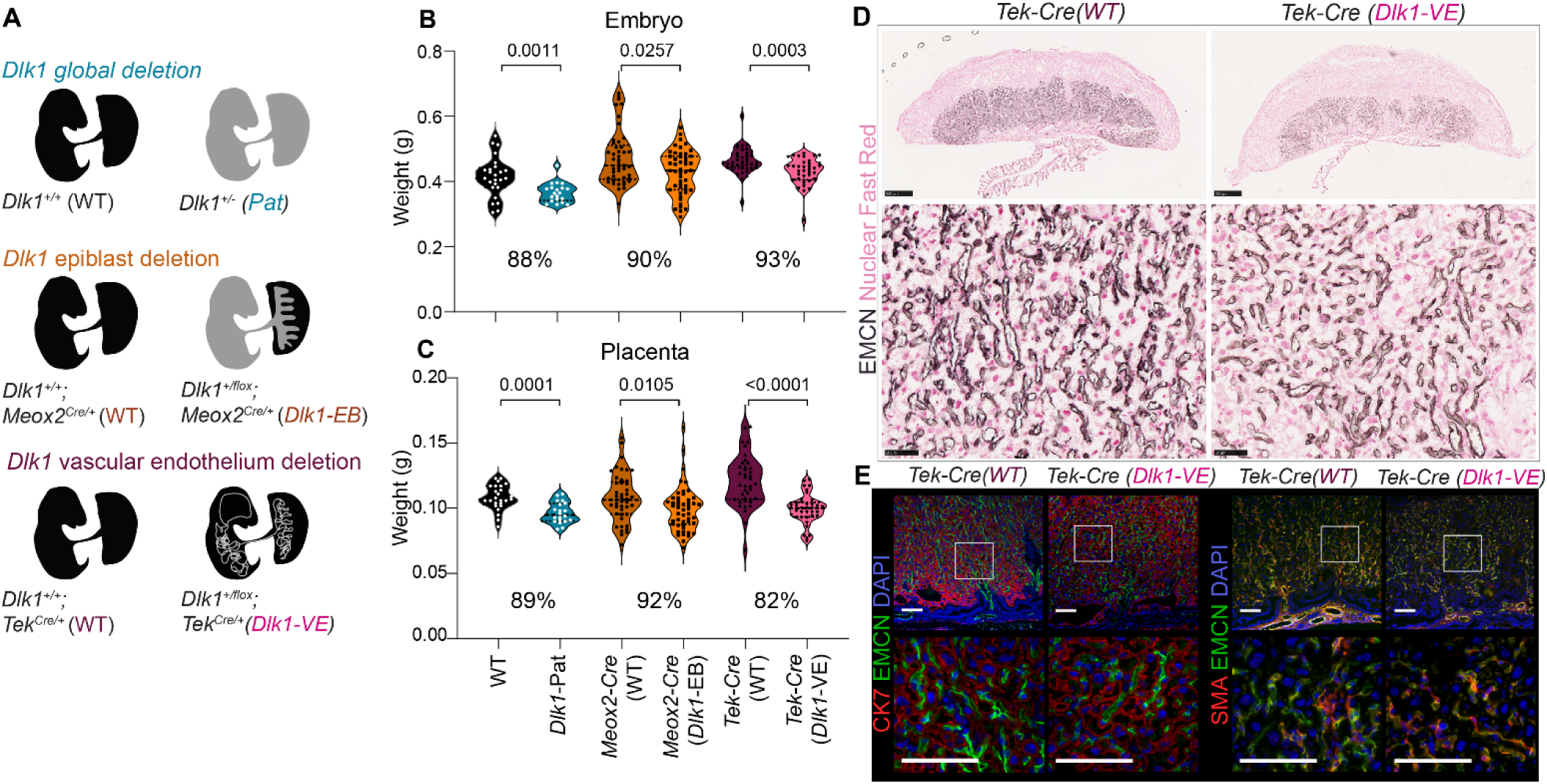
Conditional deletion of *Dlk1* in the placental endothelium causes reduced fetal-placental growth and endothelial density. A) Schematic detailing the genetic models used to explore the effect of *Dlk1*-deletion on placental and embryonic growth. Top: global deletion of *Dlk1* from all cells as a result of paternal transmission of a constitutively deleted allele. Middle: Epiblast specific deletion of *Dlk1* generated by crossing the *Meox2*-Cre with a conditional *Dlk1* allele (*Dlk1*^flox^). Bottom: vascular endothelial deletion of *Dlk1* generated by crossing the *Tek*-Cre with a conditional *Dlk1* allele (*Dlk1*^flox^). Grey areas in the embryo/placenta figures indicate the areas where *Dlk1*-deleted cells are located. B) Violin plots showing embryo mass at e15.5 in crosses shown in A. Points denote individual embryos, sexes are combined. % of corresponding WT mass is shown below the comparison. Global WT n = 25, *Dlk1*-Pat n = 20, 5 litters; *Meox2*-Cre (WT) n = 45, *Meox2*-Cre (*Dlk1*-EB) n = 48, 15 litters, *Tek*-Cre (WT) n = 40, *Tek*-Cre (*Dlk1*-VE) n = 37, 9 litters. For each cross, genotypes were compared using a Mann-Whitney U test (two-tailed) since in some cases the variances were not equal between groups. Global WT vs *Dlk1*-Pat, p = 0.0004; *Meox2*-Cre (WT) vs *Meox2*-Cre (*Dlk1*-EB), p = 0.0257; *Tek*-Cre (WT) vs *Tek*-Cre (*Dlk1*-VE), p = 0.0003. C) Violin plots showing placenta mass at e15.5 in crosses shown in A. Points denote individual placentas, n as in B. % of corresponding WT mass is shown below the comparison. For each cross, genotypes were compared using a Mann-Whitney U test (two-tailed) since in some cases the variances were not equal between groups. Global WT vs *Dlk1*-Pat, p = 0.0001; *Meox2*-Cre (WT) vs *Meox2*-Cre (*Dlk1*-EB), p = 0.0105; *Tek*-Cre (WT) vs *Tek*-Cre (*Dlk1*-VE), p < 0.0001. D) Immunohistochemistry for EMCN at e15.5 in sagittal-sectioned mouse placentas derived from *Tek*-Cre (WT) and *Tek*-Cre (*Dlk1*-VE). Positive staining is black and counterstained with nuclear fast red (pink). Bottom panels show labyrinth region of the placenta. Scale bar shows 500um in the top panels and 50 μm in the bottom panels. E) Immunofluorescence staining of *Tek*-Cre (WT) (left) and *Tek*-Cre (*Dlk1*-VE) (right) sagittal-sectioned mouse placentas at e15.5 co-stained with CK7 (pink) and EMCN (green). Images below show a high magnification view of the boxed section. For all images the scale bar = 100 µm. Images are counterstained with DAPI (blue). F) Immunofluorescence staining of *Tek*-Cre (WT) (left) and *Tek*-Cre (*Dlk1*-VE) (right) sagittal-sectioned mouse placentas at e15.5 co-stained with SMA (pink) and EMCN (green). Images below show a high magnification view of the boxed section. For all images the scale bar = 100 µm. Images are counterstained with DAPI (blue).

### Loss of Dlk1 in extraembryonic mesoderm-derived cells causes dysregulation of the Foxo1 pathway

To further characterise the molecular changes associated with *Dlk1* deletion in the placenta, we designed an experiment to compare the transcriptomes of the WT and *Dlk1*-deleted placentas at e13.5. We used a genetic labelling – flow cytometry methodology to first separate the *Dlk1*-expressing extraembryonic mesoderm population from the *Dlk1*-non-expressing trophoblast cells (Supplementary Figure 5A). We crossed *Meox2*-Cre dams with *Dlk1* heterozygous studs which were also homozygous for the dual reporter line *Rosa^mTmG^* (26) (Figure 5A, Supplementary Figure 5A). Since *Meox2*-Cre is expressed only in the inner cell mass, this resulted in offspring where the trophoblast was genetically labelled with a membrane-localised red fluorescent protein (TOM), whereas the extraembryonic mesoderm expressed GFP. This allowed us to use flow cytometry to separate the two populations of cells, which were then used for RNA sequencing (Supplementary Figure 5B).

**Figure 5.**
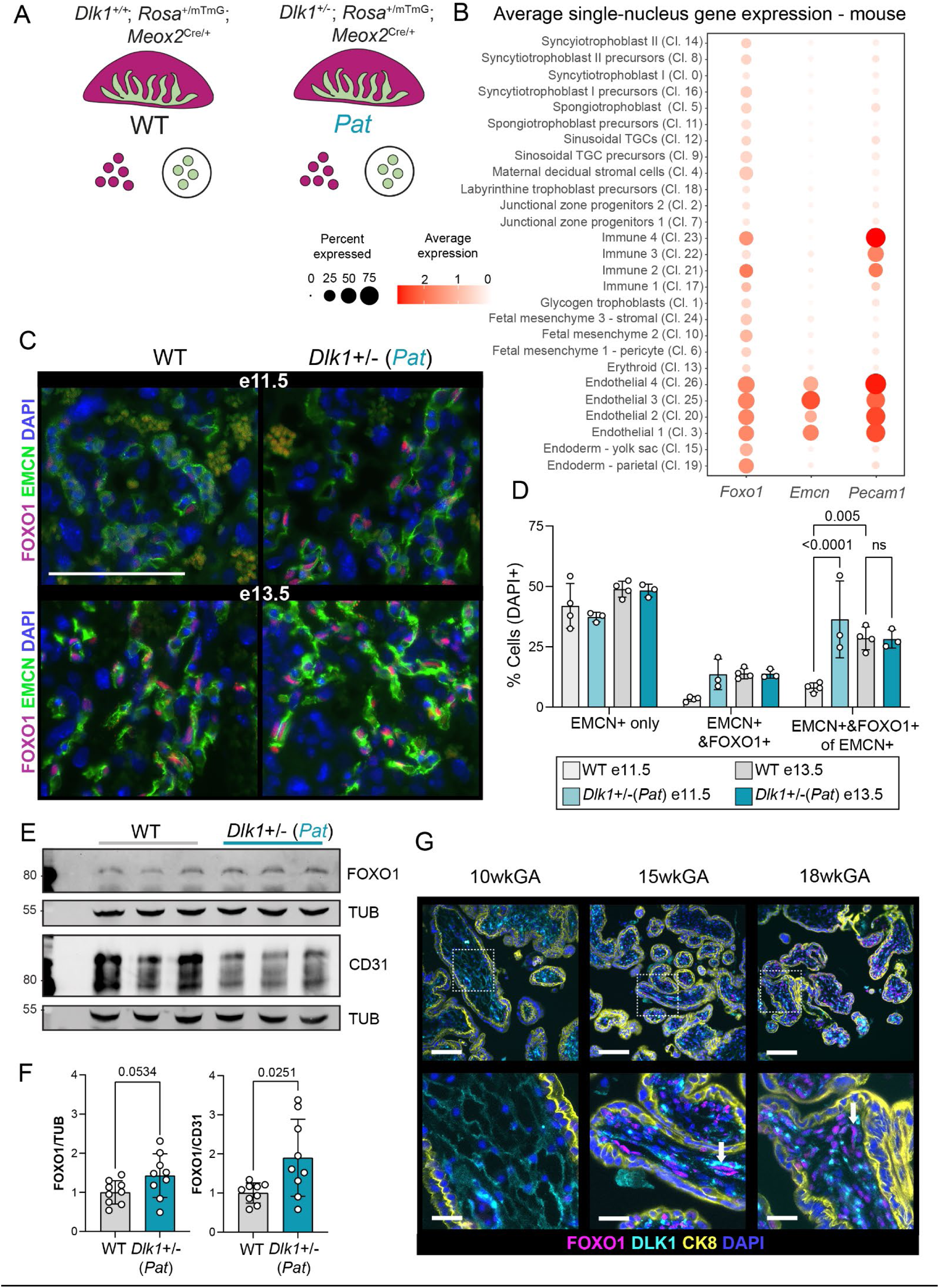
Deletion of *Dlk1* increases nuclear localization of FOXO1 in endothelial cells. A) Schematic of the experimental design. Litters of mice containing placentas with normal (WT) and ablated *Dlk1* (Pat) were generated by crossing stud *Dlk1^−/+^* Mat mice on a homozygous dual reporter Rosa^mTmG^ background to a dam heterozygous for the *Meox2*-Cre. GFP-positive cells were isolated, sorted and sequenced as described in Supplementary Figure 5A. B) Dot plot showing average single-nuclear expression of *Foxo1* across cell clusters in the mouse placenta (15), in comparison to the endothelial cell markers *Emcn* and *Pecam-1* (CD31). The color of each dot represents the average log-scaled expression of each gene across all cells in a given cluster, and the size of the dot represents the fraction of nuclei in the cluster in which transcripts for that gene were detected. C) Immunofluorescence staining of WT (left) and *Dlk1*-Pat (right) sagittal-sectioned mouse placentas co-stained with EMCN (green) and FOXO1 (pink), at e11.5 and e13.5. Nuclear localization of FOXO1 is clearly seen by overlap with DAPI (blue). For all images the scale bar = 100 µm. D) Quantification of co-localisation of nuclear FOXO1 and EMCN from images such as shown in C. N = 3-4 litter matched placentas per timepoint per genotype. Groups were compared by 2-Way ANOVA followed by Sidak’s post-hoc test comparing genotype-timepoint within each staining group. P values are shown above the bars, which indicate mean +/−SD. E) Representative Western blot of lysates from e13.5 WT and *Dlk1*-Pat placentas, probed for FOXO1 and CD31, with alpha-tubulin (TUB) used as a loading control. F) Quantification of Western blotting experiments as in E. N = 9 placentas/genotype from 3 independent litters containing 3 WT and 3 *Dlk1*-Pat placentas each. Groups are compared by Mann-Whitney U test, p values shown above the bars. G) Immunofluorescence staining of human placental villi for FOXO1 (pink) at 10 (left), 15 (middle) and 18 (right) weeks gestational age (wkGA) co-stained with DLK1 (cyan) and cytokeratin 8 (CK8, yellow). All sections are counterstained with DAPI (blue). Arrows indicate fetal blood vessels co-stained for nuclear FOXO1 and DLK1. Scale bar = 100 µm (top) and 25 µm (bottom).

To appreciate the overall differences in expression patterns between GFP and TOM populations, we performed principal component analysis (PCA) on all WT samples using the 500 genes with the largest coefficients of variation based on normalized counts. Principal component 1 (PC1) and principal component 2 (PC2) accounted for approximately 94% of all variation of the original data. To further explore and better depict the major sources of variation, all WT samples were plotted in a two-dimensional space consisting of PC1 and PC2. Samples of the GFP group and TOM group segregate from each other along PC1, demonstrating distinct expression signatures, whereas male and female samples separate along PC2 (Supplementary Figure 5C). Of note, differential gene expression analysis revealed that such sexual dimorphism originates mostly from genes located on sex chromosomes (Supporting Table 1). 4991 genes were significantly up-regulated and 3115 significantly down-regulated in GFP *vs* TOM cells from WT placentas (Supporting Table 2). We generated a heatmap of expression levels through unsupervised hierarchical clustering using the differentially expressed genes (DEGs) and found, as expected, that biological replicates of GFP and TOM populations were grouped together (Supplementary Figure 5D). Therefore, these analyses demonstrate a clear dichotomy of gene expression profiles between GFP and TOM cell populations.

We then sought to identify whether highly expressed genes in either group are enriched in known murine cell type markers. We retrieved cell type marker IDs from the PanglaoDB database (27) and performed cell type overrepresentation analysis on genes in the upper quartile of expression in either GFP or TOM samples from WT placentas (Supplementary Figure 5E). Consistent with the expectation that cells derived from the extraembryonic mesoderm are GFP+, this group was enriched for highly expressed genes encoding markers of vascular and mural cell types. Placental cell markers were the most overrepresented among highly expressed genes in the TOM group, consistent with TOM cells being of placental trophoblast origin. In addition, the mRNA abundance of well-known endothelial and stromal cell markers confirmed an extraembryonic mesoderm identity for GFP samples, while TOM samples showed higher expression of markers of placental trophoblast subpopulations (28) (Supplementary Figure 5F, data from Supporting Table 2). However, it should be noted that *Synb*, a marker of syncytiotrophoblast layer 2 (SynTII), was not detected in either group. Similarly, markers of parietal trophoblast giant cells (P-TGC) *Prl3d* and *Prl7a* were not detected across all samples. Therefore, trophoblast SynTII and P-TGC populations are likely not captured by our approach. Taken together, these observations show that GFP populations are enriched in fetal endothelial and mesenchymal cells, whereas TOM populations are enriched in trophoblast cells.

To characterize the effect of paternal knockout of *Dlk1* specifically in the extraembryonic mesoderm, we compared the transcriptomes of GFP+ populations obtained from WT and *Dlk1*-Pat animals at e13.5 (Figure 5A). We identified 477 genes significantly up-regulated in *Dlk1*-Pat compared with WT animals, and 94 down-regulated genes (Supporting Table 3). Multiple gene ontology (GO) terms were found enriched among DEGs (Supporting Table 4). These included terms related to development and growth, homeostasis of number of cells, and cellular responses to peptide hormone stimulus. KEGG pathway analysis indicated that the forkhead transcription factor (*Foxo1)* signaling pathway was enriched (Supporting Data 5). FOXO1 is a well-known modulator of vascular endothelial development, integrating the action of tyrosine kinase receptors to regulate apoptosis and vascular remodelling (29). Deletion or constitutive activation of FOXO1 in endothelial cells causes midgestation lethality with vascular defects in extraembryonic tissues (30). Examination of sn-RNAseq data from the mouse placenta (15) indicated that *Foxo1* expression is enriched in the fetal endothelial cell clusters (Figure 5B). This was confirmed by immunofluorescence staining of the placenta since FOXO1 expression overlapped predominantly with EMCN+ cells (Figure 5C). Nuclear FOXO1 was detectable in 8% of EMCN+ cells at e11.5, rising to 28% at e13.5 in the WT labyrinth, indicating that FOXO1-mediated transcription is activated in the placental fetal endothelium. In contrast, at e11.5 ∼36% of EMCN+ cells in the *Dlk1*-Pat placenta had nuclear localized FOXO1 and this level remained high at e13.5 (Figure 5D). Moreover, the overall level of FOXO1 protein increased as a proportion of the endothelial population at e13.5 (Figure 5E, F). Consistent with increased activation of FOXO1, a direct target in endothelial tip cells, endothelial cell-specific molecule 1 (*Esm1)* (31), was upregulated in the *Dlk1*-Pat GFP+ cell population (Supporting Table 3). Taken together these data indicate that failure of labyrinthine development in placentas lacking *Dlk1* is preceded by overactivation of the FOXO1 pathway in endothelial cells. In the human placenta, nuclear FOXO1 is enriched in endothelial and stromal cells of the villi, in DLK1-positive cells. FOXO1-positive nuclei appeared to be more abundant in cells at the villous tips, and to increase with gestational age (Figure 5G), as in the mouse.

### Deletion of Dlk1 increases placental hormone production

We next compared the transcriptomes of TOM populations from WT and *Dlk1*-Pat placentas (Figure 6A) and obtained 30 genes significantly up-regulated in *Dlk1*-Pat cells compared with WT cells and 20 genes down-regulated (Supporting Table 6). GO term enrichment analysis on DEGs (Supporting Table 7) revealed “prolactin receptor binding” (GO:0005148; adjusted P-value = 6.08e-05) to be the most significantly enriched term, with prolactin genes *Prl8a1*, *Prl8a8*, *Prl3b1* and *Prl2c3* being significantly more expressed in *Dlk1*-Pat compared with WT samples. Additionally, genes encoding two of the most abundant pregnancy-specific glycoproteins, *Psg16* and *Psg23*, were upregulated following *Dlk1* deletion (Figure 6B, data from Supporting Table 6). *Prl3b1* encodes Placental lactogen 2 (the homologue of human chorionic gonadotrophin 2 (CSH2)) abundantly secreted into the maternal circulation in the latter half of pregnancy, and is thought to promote maternal metabolic adaptations to pregnancy and lactation (28).

**Figure 6.**
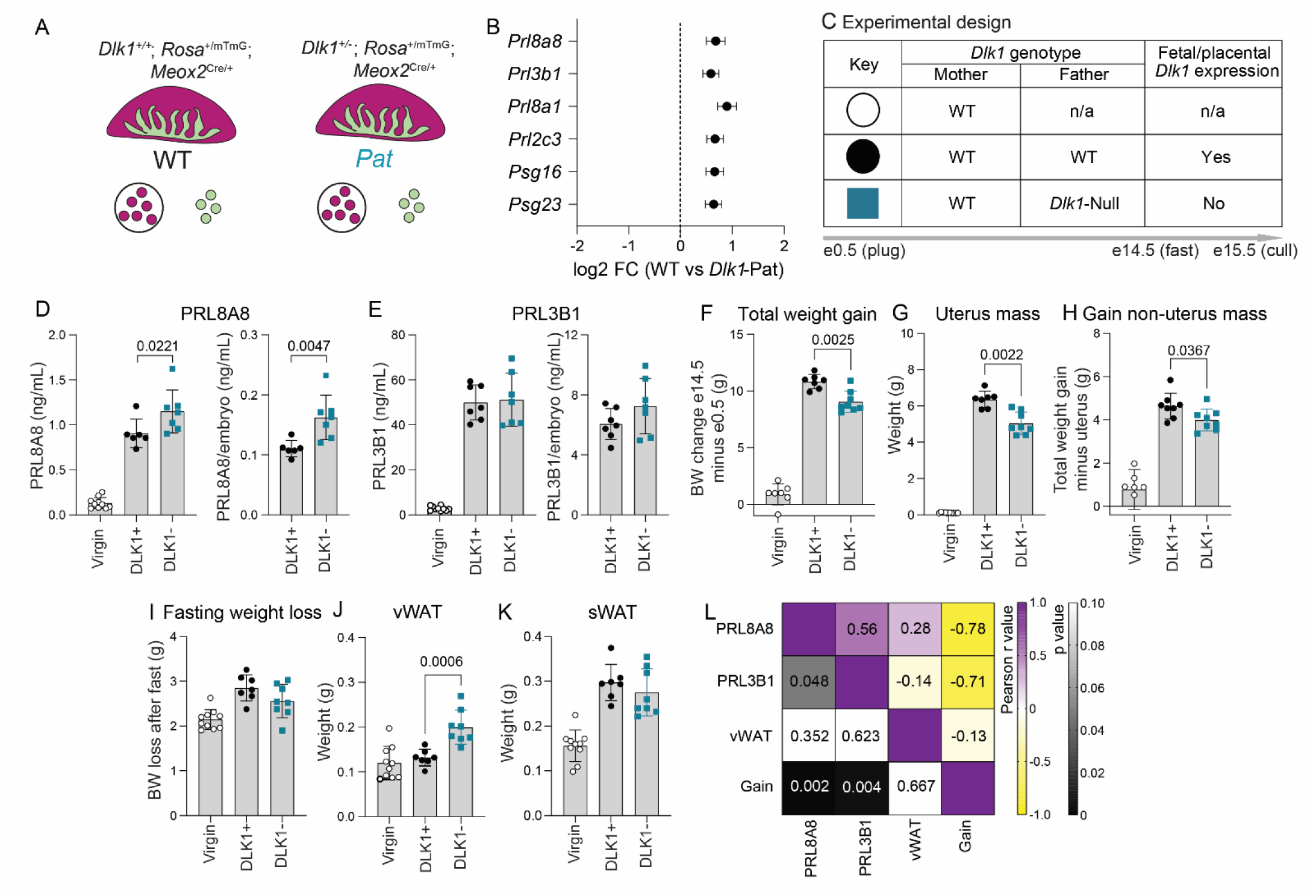
DLK1 modulates placental hormone production and maternal metabolic adaptations to pregnancy. A. Schematic of the experimental design. Litters of mice containing placentas with normal (WT) and ablated *Dlk1* (Pat) were generated by crossing stud *Dlk1^−/+^*Mat mice on a homozygous dual reporter Rosa^mTmG^ background to a dam heterozygous for the *Meox2*-Cre. TOM-positive cells were isolated, sorted and sequenced as described in Supplementary Figure 5A. B. 6/30 of the genes significantly upregulated in the *Dlk1*-Pat trophoblast cell population encoded placental hormones, including placental lactogens (*Prl*) *8a8*, *3b1*, *8a1* and *2c3* and pregnancy-specific glycoproteins (*Psg*) *16* and *23*. Differential expression (from Supporting Table 6, FDR p < 0.05) shown as fold-change increase in *Dlk1*-Pat TOM+ cells compared to WT TOM+ cells. C. Schema of the experimental design used to determine if elevated hormone-encoding gene expression in *Dlk1*-Pat placentas resulted in increased levels of hormone in the maternal circulation. 3 groups were generated, virgin WT female age-matched to the mated females, WT females mated to WT males, and WT females mated to *Dlk1^−/−^*Null males. Body weights were recorded on the day of the plug (e0.5), then on pregnancy day 14 (e14.5) immediately prior to food withdrawal at 6pm. Females were sacrificed between 8-10am on day 15 (14-16hr fast), maternal and fetal weights and samples were collected. D. Level of PRL8A8 in maternal plasma at e15.5, as total level (left) and divided by the number of embryos in the litter (right). Bars show mean +/− SD for each group, with points representing individual mice (n = 10, 6, 7). The two pregnant groups were compared by Mann-Whitney U test with significance value as indicated above the bars. E. Level of PRL3B1 in maternal plasma at e15.5, as total level (left) and divided by the number of embryos in the litter (right). Bars show mean +/− SD for each group, with points representing individual mice (n = 10, 7, 7). The two pregnant groups were compared by Mann-Whitney U test and were not significantly different. F. Body weight gained per female determined by subtracting the mass recorded at e0.5 from the mass at e14.5. The two pregnant groups were compared by Mann-Whitney U test with significance value as indicated above the bars. G. Uterus mass at e15.5 for the 3 groups of mice, compared as shown in F. H. Body weight gained per female minus the uterine weight, compared as shown in F. I. Females were fasted for 14-16hr overnight on the 14^th^ day of pregnancy, and weight change in this interval was recorded. The two pregnant groups were compared by Mann-Whitney U test and were not significantly different. J. Mass of the virgin or pregnant female visceral adipose tissue (vWAT, including the perigonadal and perirenal white adipose depot) post-sacrifice at e15.5, compared as in F. K. Mass of the virgin or pregnant female subcutaneous adipose tissue (sWAT, including the posterior dorsal and ventral subcutaneous depots) post-sacrifice at e15.5, compared as in F. L. Correlation matrix of Pearson correlations of hormone levels (PRL8A8 and PRL3B1), vWAT and weight gain (Gain, non-uterus mass) for all pregnant females including the DLK1+ and DLK1- groups (n = 13-14). Positive Pearson’s correlation values are shown in purple and negative correlations in yellow, with value shown inside the grid. Corresponding p values are shown in the bottom left of the grid, not corrected for multiple comparisons.

We generated an additional cohort of mice in which to test if elevated gene expression of placental lactogens in the *Dlk1*-Pat trophoblast cells resulted in elevated maternal hormone levels and related alterations to maternal metabolic indices (Figure 6C). Here we generated two groups of mice, one where the mother and all placentas/embryos contained a WT copy of *Dlk1* (DLK1+ litters), the other where the mother was WT but all of her offspring had received a *Dlk1*-knockout allele from their father, hence were each *Dlk1*-Pats (DLK1- litters). In the latter group all of the placentas should have elevated prolactin gene expression. We sacrificed the mother at e15.5 then collected maternal plasma for hormone measurements. A third group of age-matched virgin WT females were collected in parallel. As expected, circulating levels of PRL3B1 and PRL8A8 were considerably elevated by pregnancy (Figure 6D, E). PRL8A8 hormone levels were increased (both total levels and levels per pup) in mothers carrying DLK1- litters (DLK1- mothers) compared to mothers carrying DLK1+ litters (DLK1+ mothers) in a similar degree to the increase in placental gene expression (Figure 6D, ∼30% and 50% increased, respectively). PRL3B1 hormone levels were not different between the two groups (Figure 6E).

We previously reported that maternal pregnancy weight gain and adipose mass is influenced by *Dlk1* levels in the conceptus, especially following maternal fasting (9). Consistently, here we found that mothers in DLK1- pregnancies gained less body mass over the pregnancy period compared to DLK1+ mothers (Figure 6F). This was firstly due to a decrease in uterine mass (Figure 6G), which was reduced in DLK1- mothers because of the expected reduction in embryonic weight and a smaller litter size (Supporting Data for Figure 6). Secondly, the non-uterine mass of the DLK1- mothers increased less over the pregnancy period compared to DLK1+ mothers (4.0g vs 4.6g respectively, Figure 6H).

Following an overnight fast, DLK1- mothers lost less weight and retained more visceral white adipose tissue (vWAT), but not subcutaneous white adipose tissue (sWAT), compared to DLK1+ mothers (Figure 6I-K). Conceptus-derived DLK1 therefore increases net weight gain in maternal tissues over the course of pregnancy and promotes the liberation of energy from visceral adipose tissue stores following fasting. To understand if either of these actions are related to the production of lactogens from the placenta, we asked if maternal circulating hormone levels were correlated with weight gain or adipose mass. We found no relationship between circulating PRL8A8 or PRL3B1 with adipose mass. In contrast there was a strong negative correlation between levels of these hormones and maternal weight gain (Pearson’s R = −0.78 for PRL8A8 and −0.71 for PRL3B1, p = 0.002 and 0.004, respectively). In addition, levels of the two hormones were significantly co-correlated (Pearson’s R = 0.56, p = 0.048) (Figure 6L). Increasing these prolactin levels may therefore inhibit maternal body weight gain in pregnancy. Taken together these data indicate that the maternal body mass-promoting action of DLK1 in pregnancy may be mediated by the production of lactogens by placental trophoblast cells.

## Discussion

*Dlk1* is highly expressed during embryogenesis and its levels are linked to intrauterine growth. Here we confirm that placental *Dlk1* expression is limited to the extraembryonic mesoderm (EEM) lineage of the developing placenta, and the gene is not expressed in trophoblast cells (32, 33). In human placentas, the expression of *DLK1* is less well defined, with some reports of trophoblast expression (13). We used data from single-nucleus sequencing studies of the human placenta (19), as well as immunohistochemistry techniques, to show that, as in the mouse, DLK1 expression is limited to the EEM. Since DLK1 is a secreted molecule, previous studies may have detected EEM-derived circulating DLK1 associated with trophoblast cells. Establishing the developmental origin of this signalling molecule is important, since this information is essential to discover the mechanism of its normal role in growth.

Since *Dlk1* is expressed widely in embryonic tissues as well as in the placenta, we sought to understand to what extent the growth restriction phenotype following *Dlk1* deletion was caused by a role of this gene in the placenta. We found that *Dlk1* deletion resulted in the failure of the placental labyrinth to expand beyond embryonic day e13.5. This placental region is the site of gas and nutrient exchange between mother and fetus, therefore reduction in this compartment could result in failure to support fetal growth. Using conditional targeting we deleted *Dlk1* in the endothelial cells of the labyrinth. This caused a similar placental mass reduction to the *Dlk1* global deletion and accounted for around half of the reduction in embryonic mass. We concluded that placental endothelial *Dlk1* production is required for the embryo to reach its full growth potential.

Our conditional deletion experiments and subsequent transcriptomics analysis showed that *Dlk1* acts cell autonomously in endothelial cells to alter the density of the vascular network. Angiogenesis of the placental fetal vasculature is driven by vascular endothelial growth factor (VEGF) signalling (34). In the mouse, VEGFA is produced by the labyrinthine syncytiotrophoblast cells, and acts on VEGF receptors (encoded by *Flt1* and *Kdr*) on endothelial cells (15, 35, 36). Global or endothelial cell-specific deletion of the gene encoding *Foxo1* results in midgestation lethality following failure of yolk-sac and placental angiogenesis. *Foxo1*-deficient ECs fail to respond to angiogenic stimuli, including VEGF (37, 38). Using a retinal angiogenesis model, Wilhelm et. al (30) elegantly showed that FOXO1 is expressed in the tip cells of the plexus, where vessels remodel and proliferation is supressed. Deletion of *Foxo1* in this system resulted in failure of angiogenic sprouting and in a hyperplastic vascular network. Expression of constitutively active nuclear-localised FOXO1 in endothelial cells also caused mid-gestation lethality, and in retinal assays this produced a sparse vascular network with reduced EC diameter. In the placenta we observed nuclear FOXO1 localisation commencing at e11.5 of development in a small proportion of ECs (8%). By e13.5, 28% of ECs expressed nuclear FOXO1 in the WT placenta (Figure 5). The increase in FOXO1 activation is concurrent with the increase in endothelial branching and vessel elaboration of the placenta, as can be seen by the marked increase in EC density and reduced cross sectional area between e11.5 and e13.5 (Figure 3). *Dlk1*-deleted placentas exhibit reduced endothelial density and premature terminal vessel maturation, and this is accompanied by early nuclear localisation of FOXO1 and upregulation of the tip cell marker *Esm1*. We propose that *Dlk1* modulates the timing of endothelial branching morphogenesis by regulating VEGF sensitivity via expression of FOXO1. *Dlk1* has previously been shown to negatively regulate angiogenesis in cultured ECs by interaction with the Notch signalling pathway (39). The phenotype of the *Dlk1*-deleted placental vasculature is not consistent with this role, and we did not observe any changes to the expression of *Notch1* or its downstream transcriptional mediator *Hes1* (Supplementary table). However, the timing of vessel sprouting and maturation is highly dynamic, as is the response of ECs to VEGFs (31). Our data could be reconciled with the previous study if these timing differences are taken into account. We note that aortic explants from *Dlk1* mutant mice showed increased sprouting in the Rodriguez study (39), which is consistent with our placental data.

VEGF signalling is a key player in human placental development. From the end of the first to the beginning of the second trimester, immature placental villi comprising a trophoblast outer layer and a stromal core become increasingly vascularised and elaborated. Extra-embryonic mesoderm-derived endothelial cells grow, branch and form close contact with the trophoblast layer maximising maternal-fetal exchange (34). Failure of this process is associated with fetal growth restriction (40). In the first trimester, VEGF is produced by the cytotrophoblast, and as pregnancy continues production from Hoffbauer and mesenchymal stromal cells contributes to this signal. As in rodents, VEGF receptors are present on endothelial cells and they respond in the early phase of pregnancy by undergoing branching angiogenesis to define the vascular tree (34). In later gestation the actions of placental growth factor (PlGF) and angiopoietin signals act to expand and refine the vascular network, resulting in terminal villi that comprise the interhemal barrier (34). In the first trimester (prior to 12wkGA) we did not observe any nuclear FOXO1 staining, consistent with the lack of vascularised secondary villi at this timepoint. We observed FOXO1 expression in the human placental villi from 12wkGA (not shown), with robust levels in nuclei of endothelial and stromal cells of the second trimester villi. Activation of FOXO1 in the human placenta is therefore coincident with extensive angiogenic remodelling. As in the mouse placenta, extraembryonic mesoderm-derived cells with nuclear FOXO1 were also DLK1-positive, suggesting a commonality to this mechanism across species.

Whilst *Dlk1* expression is confined to the EEM, we unexpectedly observed differential gene expression in sorted trophoblast cells between *Dlk1*-deleted and WT placentas. A small number of genes affected were upregulated in the *Dlk1*-deleted trophoblasts and encoded abundantly expressed placental hormones. We speculated that this increase in hormone-producing mRNA could result in elevated levels of their products in the maternal circulation and thereby influence maternal metabolic adaptations to pregnancy. Since we have already linked low circulating DLK1 to alterations in maternal energy homeostasis (9), we investigated hormone levels and maternal metabolic indices in a further cohort of pregnant mice. Here we compared mothers where the whole litter had a deleted *Dlk1*-gene, therefore all of the placentas had elevated hormonal mRNA production, with WT-only litters (controls). As predicted, one hormone, PRL8A8, had elevated levels in the maternal circulation of *Dlk1*-deleted litters. Levels of PRL8A8 and another abundant hormone, PRL3A1, were negatively correlated with maternal weight gain, in concert to the effect of DLK1. We speculate that the hormone-producing cells of the trophoblast may be mediating some of the actions of DLK1 on maternal energy homeostasis. There is a growing body of evidence that placental endocrine production is controlled by imprinted gene expression dosage (41, 42). Furthermore, altered expression of other imprinted genes in *Dlk1*-deleted placentas (Supporting Table 3) lends further evidence to the existence of an imprinted gene network which co-ordinately regulates growth (43). Variants at imprinted loci are overrepresented in genome-wide association studies of birthweight (44), and their actions on physiology more widely have likely been underestimated (45).

Taken together, here we provide mechanistic details describing how loss of *Dlk1* in the placenta causes fetal growth restriction in the mouse. We propose that low DLK1 in endothelial cells of the placenta alters the rate of branching morphogenesis, by altering nuclear levels of the transcription factor FOXO1 and hence sensitivity to angiogenic signals such as VEGF. We show that a similar activation of FOXO1 is associated with angiogenesis in the human placenta, and is coincident with local expression of DLK1. Overall, our data suggests that human growth restriction associated with DLK1 deficiency is caused by placental vascular insufficiency.

### Limitations

This work has several limitations. Firstly, the *Tek*-Cre-derived deletion of *Dlk1* in endothelial cells resulted in the upregulation of *Dlk1* in the adjacent stromal cells of the placenta. This unexpected result makes the conclusion that the effect of loss of DLK1 in the endothelium alone causes placental insufficiency secondary to the caveat that the phenotype could also be due to alterations to stromal cells or elevated circulating DLK1 in the placenta. However, since the endothelial phenotype of the global *Dlk1*-deleted placental vasculature is similar to the EC deletion, we believe the most parsimonious explanation is that the vascular phenotype is caused by loss of EC-*Dlk1* expression rather than gain of stromal expression. We measured the levels of circulating placental lactogens, PRL8A8 and PRL3B1 using commercial ELISA kits. These reagents performed as predicted with regard to performance of standards and negative controls. Moreover, both hormones were significantly elevated in pregnancy compared to in virgin plasma, as expected (Figure 6). However, in the absence of mouse lines lacking the genes encoding these hormones, we cannot be completely certain that we are accurately measuring the expression of these specific hormones and not other members of this highly conserved hormone family.

## Materials and Methods

### Sex as a biological variable

For experiments on mice both sexes of embryo and placenta were included in the study and the sexes are combined unless otherwise stated. Sexes were combined since the same influence of the global *Dlk1* deletion embryonic or placental weight was observed in both sexes (Supplementary Figure 3A&B). For transcriptomic studies we used both male and female placental samples and included sex as a random variable in the analysis. Specifically, RNA-seq gene counts were normalized using the DESeq2 R/Bioconductor package (v. 1.28.1(46)) accounting for all factors (e.g. sex, litter, genotype) that may affect expression of each gene, followed by a regularized log transformation.

### Statistics

All statistical tests were performed using the GraphPad Prism Software version 9 for Windows, GraphPad Software, San Diego California USA, www.graphpad.com. Specific tests, significance values and number of samples analysed are indicated in the respective figure/table legends, and all error bars represent the standard deviation (SD). Data points in graphs represent individual animals as biological replicates. Outliers were removed using Grubb’s test within the Prism application. Datasets were checked for normality, and non-parametric analysis performed if this property could not be confirmed.

### Study approval

All animal procedures were carried out in accordance with the recommendations provided in the Animals (Scientific procedures) Act 1986 of the UK Government, under UK Home Office licence PP2573661. Paraffin-embedded human placental material was obtained from the MRC-Wellcome Trust Human Developmental Biology Resource (HDBR), under their project ethical approvals; IRAS project ID 326492.

### Data availability

Transcriptomics data are available in GEO (GSE313753). Further information and source data can be found in the Supplementary Tables and Supporting data values file.

## Supporting information

Combined_supplementary_data

## Author contributions

(CRediT)

**MLV** methodology, investigation, formal analysis, visualization, writing - review & editing; **RE** conceptualisation, methodology, investigation, formal analysis, visualization, data curation, writing - review & editing, **VS** methodology, investigation, formal analysis, visualization, writing - review & editing, **CS** investigation, formal analysis, **DK** investigation, **EM** investigation, **CD** investigation, **MC** conceptualisation, methodology, validation, investigation, formal analysis, visualization, supervision, writing – original draft, project administration, funding acquisition.

## Funding Support

Project funding was provided by the UK Research and Innovation grants (UKRI) grants MR/L002345/1, MR/R022836/1 and BB/X007758/1 to **MC**. **MLV** was supported by a PhD studentship from the NIHR Biomedical Research Centre at Guy’s and St Thomas’ NHS Foundation trust and King’s College London. **CS** was supported by the Erasmus+ programme.

## Acknowledgements

We thank Dr Miguel Constancia, University of Cambridge UK for sharing the *Meox2*-Cre line, and Dr Ionel Sandovici University of Cambridge UK for the placental dissociation protocol which was the starting point for our method.

## Notes

The authors have declared that no conflict of interest exists.

### Competing Interest Statement

The authors have declared no competing interest.

