## Supplementary material for "Placental insufficiency causes fetal growth restriction in mice lacking *Delta-like homologue 1*": Combined_supplementary_data

**<u>Supplementary information</u>**

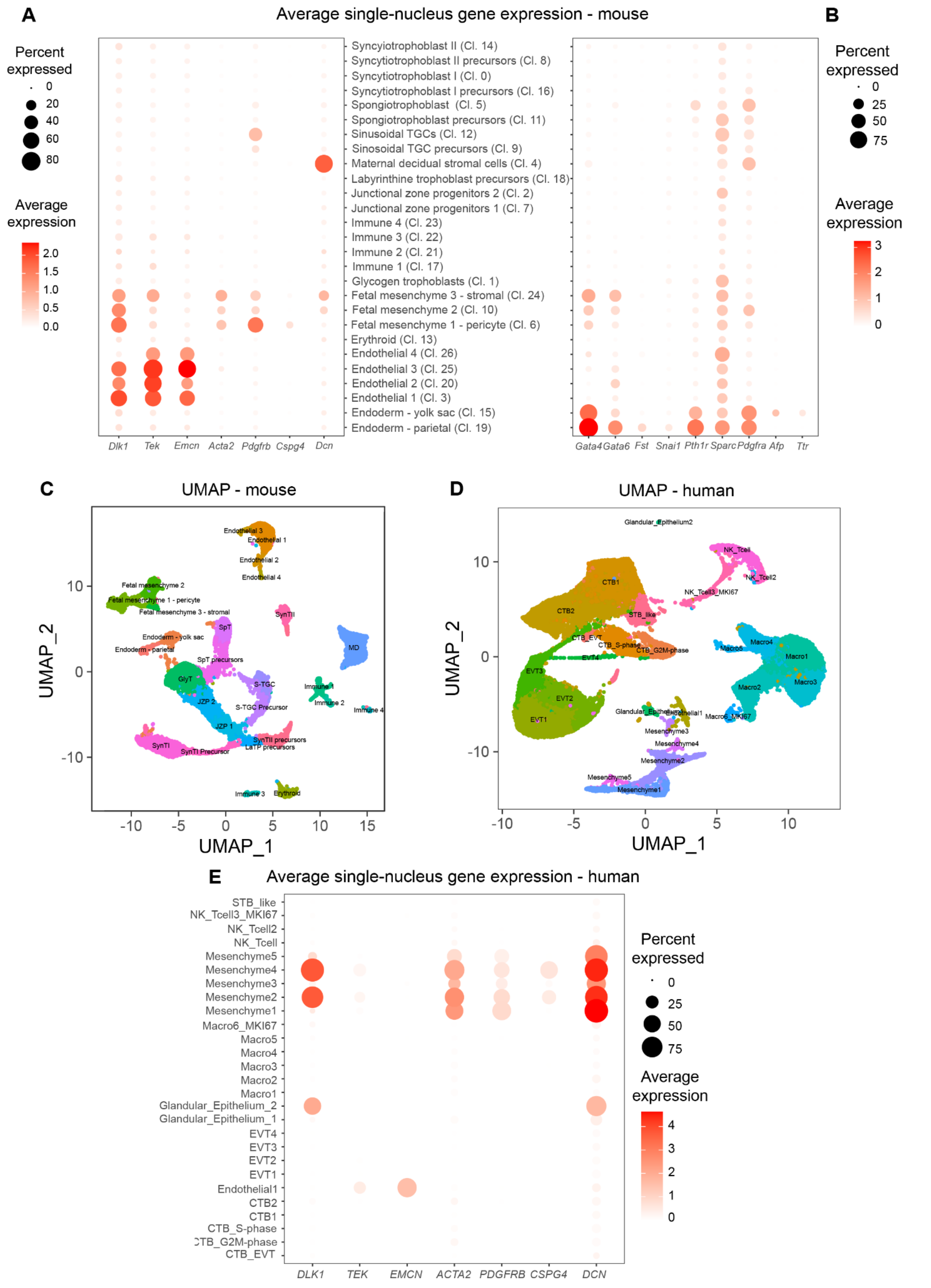


**Supplementary Figure 1. Expression of cell-type specific markers in sn-RNAseq clusters.**

A) Dot plot showing average single-nuclear expression of *Dlk1* across cell clusters in the mouse placenta(1), in reference to other endothelial cell (*Tek*, *Emcn*) and mesenchymal cell (*Acta2*, *Pdgfrb*, *Cspg4*, *Dcn*) markers. The color of each dot represents the average log-scaled expression of each gene across all cells in a given cluster, and the size of the dot represents the fraction of nuclei in the cluster in which transcripts for that gene were detected. Cl. # refers to the original cluster designation in the source paper.
B) Dot plot showing average single-nuclear expression of markers relevant to visceral endoderm fate and their enrichment in clusters 15 and 19.
C) Uniform Manifold Approximation and Projection (UMAP) plot showing relationships between cell clusters in data from the mouse placenta (1).
D) Uniform Manifold Approximation and Projection (UMAP) plot showing relationships between cell clusters in data from the mouse placenta (2).
E) Dot plot from a sn-RNAseq dataset of human placenta (2), illustrating that *DLK1* is expressed in cells arising from the extraembryonic mesoderm.

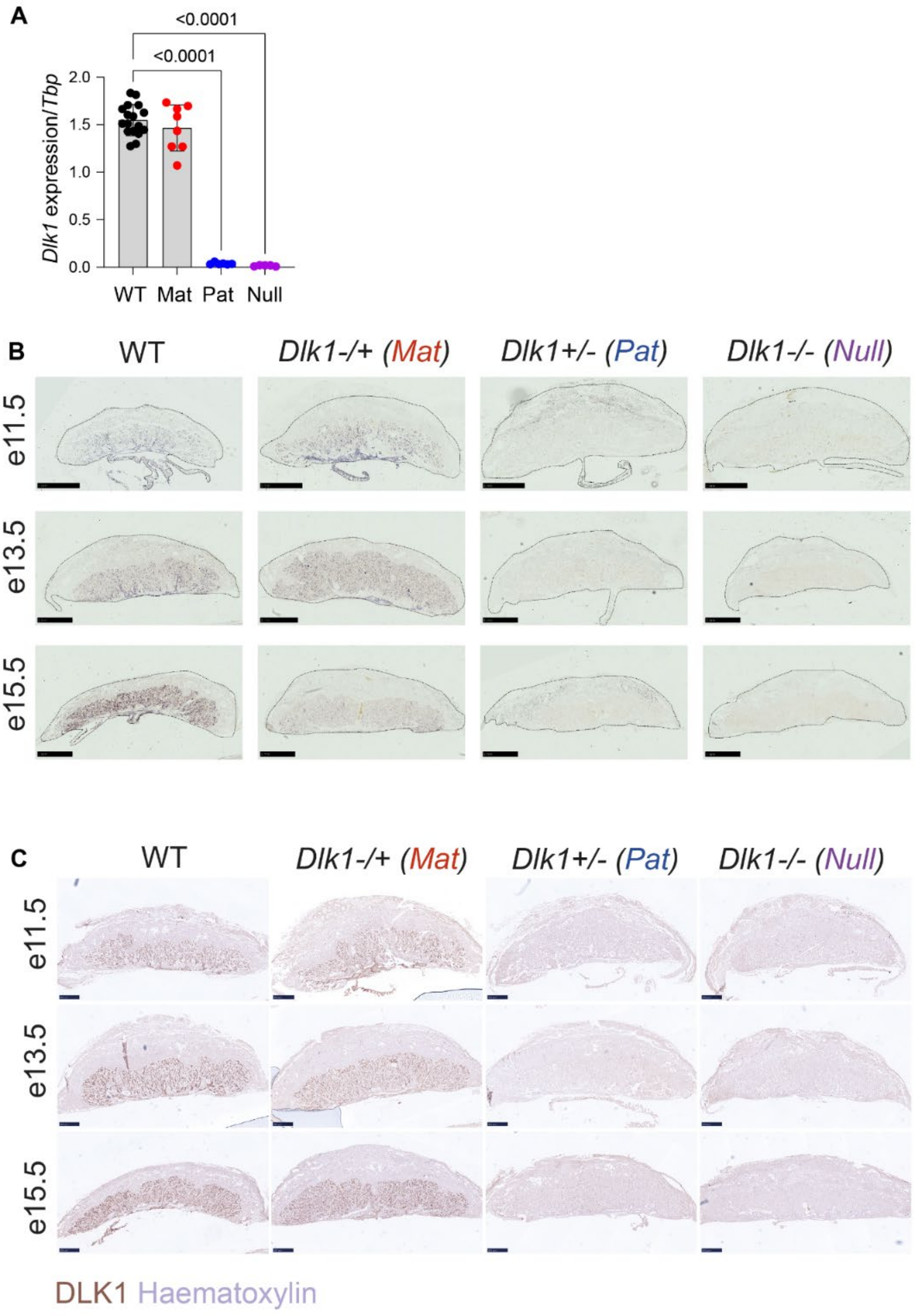


**Supplementary Figure 2. Imprinting is maintained in the placenta.**

A) Real-time quantitative PCR (RT-qPCR) for *Dlk1* in placentas from WT (*Dlk1*$^{+/+}$, n = 16), heterozygotes with the *Dlk1* deletion on the maternally-inherited allele (*Dlk1*$^{-/+}$, Mat, n = 8), heterozygotes with the *Dlk1* deletion on the paternally-inherited allele (*Dlk1*$^{+/-}$, Pat, n = 6), and homozygotes inheriting a *Dlk1*-deletion from both parents (*Dlk1*$^{-/-}$, Null, n = 5). Data is normalised to the *TATA-box binding protein* gene (*Tbp*). Relative expression is shown in a bar chart with mean +/- SD and individual data points. Genotypes were compared by One-Way ANOVA (two-tailed) with Dunnett's Multiple comparison test comparing each genotype to WT, and Pat to Null. WT vs Mat, p = 0.669, WT vs Pat, p < 0.0001***, WT vs Null, p < 0.0001***, Pat vs Null p = 0.999.

B) In-situ hybridisation for *Dlk1* at embryonic stages e11.5, e13.5 and e15.5 in sagittal-sectioned mouse placentas derived from heterozygous inter-crosses of *Dlk1*-deleted parents, or to WT. Positive staining is stained in purple and counterstained with Nuclear Fast Red. Scale bar shows 500 µm.

C) Immunohistochemistry for DLK1 at embryonic stages e11.5, e13.5 and e15.5 in sagittal-sectioned mouse placentas derived from heterozygous inter-crosses of *Dlk1*-deleted parents, or to WT. Positive staining is stained in brown and counterstained in blue with Haematoxylin. Scale bar shows 500 µm.

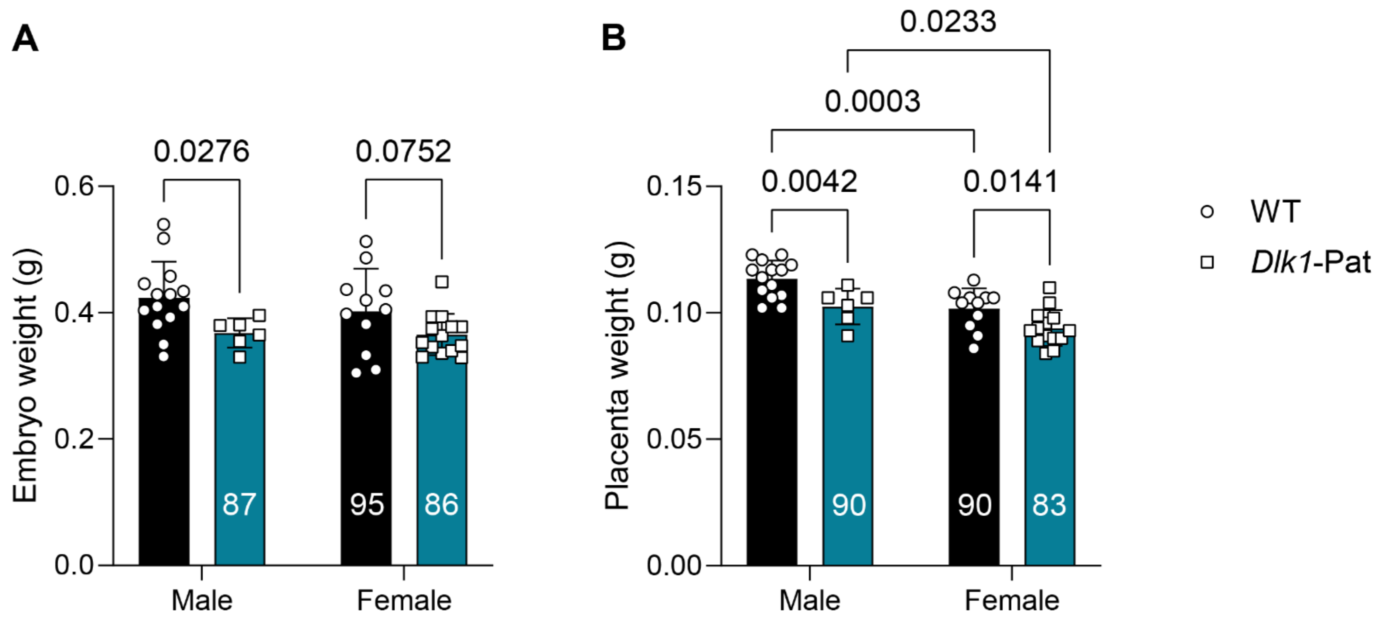


**Supplementary Figure 3. Global *Dlk1* deletion reduces embryo and placental mass in both sexes**.

A. Bar charts showing embryo mass at e15.5 following paternal transmission of the global deletion of *Dlk1*, shown in Figure 4A&B, separated by fetal sex, points denote individual embryos. % of corresponding WT male mass is shown below within the bar for comparison. Genotype and sex were compared by Two-way ANOVA; genotypes were significantly different (p = 0.006), but sexes were not (p = 0.452) nor was the interaction between them (p = 0.553). Significance p values obtained from post-hoc testing (Uncorrected Fisher's UST) are displayed above the bars. Bars show mean +/- SD.

B. Bar charts showing placenta mass at e15.5 following paternal transmission of the global deletion of *Dlk1*, shown in Figure 4A&C, separated by fetal sex, points denote individual placentas. % of corresponding WT male mass is shown below within the bar for comparison. Genotype and sex were compared by Two-way ANOVA; genotypes (p = 0.0003) and sexes (p < 0.0001) were significantly different, with no interaction between them (p = 0.485). Significance p values obtained from post-hoc testing (Uncorrected Fisher's UST) are displayed above the bars. Bars show mean +/- SD.

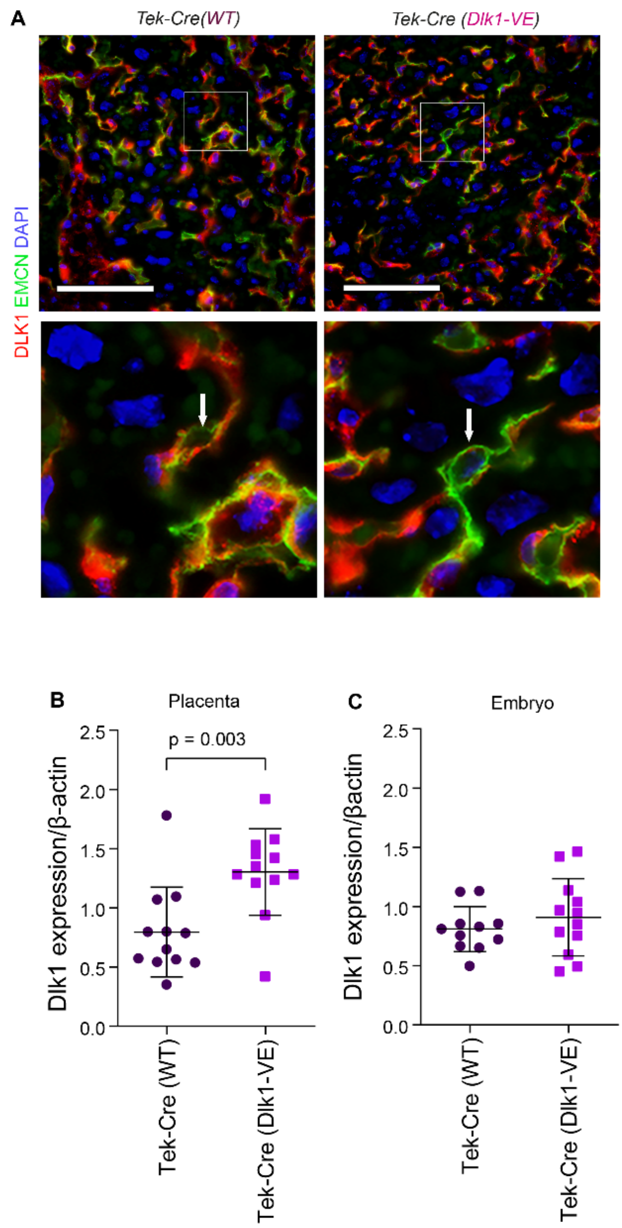


**Supplementary Figure 4. Conditional deletion of *Dlk1* using the *Tek*-Cre results in the loss of DLK1 in EC but not FM cells**.

A) Immunofluorescence staining of *Tek*-Cre (WT) (left) and *Tek*-Cre (*Dlk1*-VE) (right) sagittal-sectioned mouse placentas at e15.5 co-stained with DLK1 (pink) and EMCN (green). Images below show a high magnification view of the boxed section. For all images the scale bar = 100 µm. Images are counterstained with DAPI (blue). Arrows indicate endothelial cells.

B) RT-qPCR for *Dlk1* in placentas at e15.5 from *Tek*-Cre (WT), n = 12, and *Tek*-Cre (*Dlk1*-VE), n = 12. Data is normalised to β-actin. Relative expression is shown in a bar chart with mean +/- SD and data points representing individual placentas. Genotypes were compared by Student's t-test (two-tailed), p = 0.003.

C) RT-qPCR for *Dlk1* in embryos at e15.5 from *Tek*-Cre (WT), n = 11, and *Tek*-Cre (*Dlk1*-VE), n = 12. Data is normalised to β-actin. Relative expression is shown in a bar chart with mean +/- SD and data points representing individual placentas. Genotypes were compared by Student's t-test (two-tailed), p = 0.388.

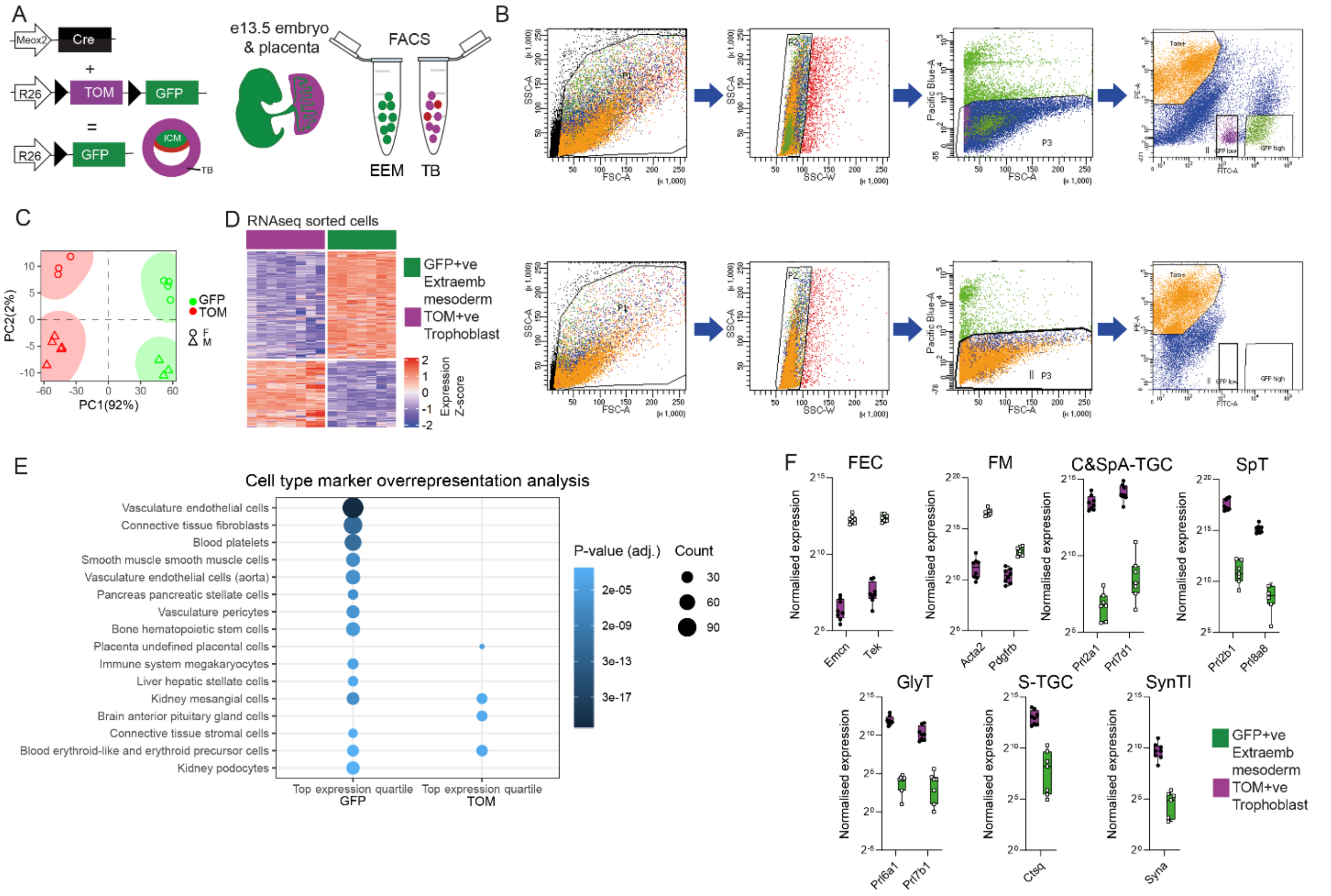

**Supplementary Figure 5. Genetic labelling and flow sorting allows separation of cells from different placental compartments.**

A) Schema of genetic crosses used to generate labelled placentas used in the RNAseq experiment. Female *Meox2*-Cre mice are crossed to stud dual reporter mice (*Rosa*$^{mTmG}$). Expression of Cre recombinase switches red fluorescent (TOM) to green fluorescent (GFP) gene expression. *Meox2* is expressed in inner cell mass (ICM) precursors of the extraembryonic mesoderm (EEM), but not the trophoblast (TB), therefore the e13.5 placenta expresses TOM in trophoblast and GFP in the endothelium and mesenchyme. Fluorescence activated cell sorting (FACS) of dissociated placentas allows separation of GFP+ and TOM+ cells for transcriptomic analysis by RNAseq.

B) Example of flow and gating strategy used to separate GFP and TOM labelled cells from the placenta. Top – e13.5 placenta of genotype *Meox2*$^{Cre/+}$; *Rosa*$^{mTmG/+}$, bottom – control e13.5 placenta which was *Meox2*-Cre negative (*Meox2*$^{+/+}$; *Rosa*$^{mTmG/+}$), therefore GFP+ cells were absent.

C) Principal component analysis (PCA) plot for GFP+ (GFP) and tdTomato+ (TOM) samples (WT only). The PCA was performed on all samples using the 500 genes that had the largest coefficients of variation based on normalized counts. GFP and TOM samples are separated along principal component 1 (PC1), accounting for 94% of the variance, and male (M) and female (F) samples are segregated along principal component 2 (PC2).

D) Heatmap of z-scored expression levels of differentially expressed genes (TOM *vs* GFP; n = 8106) with samples (columns) and genes (rows) ordered by complete linkage clustering on Euclidean distances.

E) Cell type overrepresentation analysis chart showing significant enrichment of vascular cell type markers among genes at the top quartile of expression in FACS-purified GFP+ placental cells (WT), as well as enrichment of undefined placental cell markers among highly expressed genes in TOM+ cells (all WT). Marker gene IDs were obtained from the PanglaoDB database (3). P-values were calculated by hypergeometric distribution and adjusted for multiple hypotheses testing by the Benjamini-Hochberg procedure.

F) Boxplots showing expression levels of cell markers in GFP and TOM populations (from WT placentas). FEC, fetal endothelial cells; FM, fetal mesenchyme; C-TGC, canal trophoblast giant cell; SpA-TGC, spiral artery trophoblast giant cell; GlyT, glycogen trophoblast; S-TGC, sinusoidal trophoblast giant cell; SpT, spongiotrophoblast; SynTI, syncytiotrophoblast cells.

### **Supplementary Materials and Methods**

#### Mice

##### *Husbandry*

Mice were maintained on a 12-hour light/dark cycle in a temperature and humidity-controlled room and re-housed in clean cages weekly. All mice were fed *ad libitum* and given fresh tap water daily. Weaning was performed at postnatal day (P) 21 into single-sex groups (5 per cage maximum) or occasionally singly housed. Embryos were generated through timed matings (either as a pair or a trio). Noon on the day of the vaginal plug was considered as embryonic day (e)0.5.

##### *Strains*

All mice were maintained on a C57BL6/J background. Global *Dlk1*-deleted mice [*Dlk1*$^{tm1Srba}$ , (4), hereafter *Dlk1*-Mat (*Dlk1*$^{-/+}$ ), *Dlk1-Pat* (*Dlk1*$^{+/-}$), or *Dlk1-Null* (*Dlk1*$^{-/-}$)] were maintained as a heterozygous stock of animals that inherit the deleted allele from the silent maternal copy (*Dlk1*-Mats), as described previously(5). For the flow sorting of placental cells, *Dlk1*-Null mice were crossed onto the *Rosa26*$^{mTmG}$ dual reporter line (Gt(ROSA)26Sor$^{tm4(ACTB-tdTomato,-EGFP)Luo/J}$, (6) hereafter *Rosa26*$^{mTmG}$) to generate *Dlk1*$^{-/+}$(Mat); *Rosa26*$^{mTmG/mTmG}$ sires. Conditional *Dlk1*-deleted mice (*Dlk1*$^{tm1.1Jvs}$, (7) hereafter *Dlk1*-flox (*Dlk1*$^{+/flox}$)) were crossed onto the *Rosa26*$^{mTmG}$ line to generate *Dlk1*$^{flox/+}$; *Rosa26*$^{mTmG/mTmG}$ sires. *Dlk1*-epiblast-specific deletion (*Dlk1*-EB) was generated by crossing Meox2-Cre (*Meox2*$^{tm1(cre)Sor}$, (8)) heterozygous dams to *Dlk1*$^{flox/+}$; *Rosa26*$^{mTmG/mTmG}$ sires, as described in (9). *Dlk1*-endothelial-specific deletion (*Dlk1*-VE) was generated by crossing Tek-Cre (*Tg(Tek-cre)12Flv(10)*) homozygous dams to *Dlk1*$^{flox/+}$; *Rosa26*$^{mTmG/mTmG}$ sires.

##### *Genotyping*

Genotyping and sexing PCRs for embryos were performed on ear and embryonic tail biopsies with DNA extracted using DNAReleasy (LS02, Anachem) according to the manufacturer's instructions. PCR was performed using REDTaq® ReadyMix™ PCR Reaction Mix (R2523, Merck), except for the *Dlk1*-flox genotyping which used KAPA2G Fast HotStart ReadyMix (2GFHSRMDKB, Merck). All genotyping and sexing primers can be found in Supplementary Table 1.

| Target | Forward (5'-3') | Reverse (5'-3') |
|---|---|---|
| *Dlk1* global deletion | CCAAATTGTCTATAGTCTCCC | CTGTATGAAGAGGACCAAGG<br>CATCTGCACGAGACTAGTG |
| *Dlk1*-flox | AGATTCCCCCACCTCCAAC | TTCCCAAACTGGACATGAGC |
| *Meox2*-Cre | GGACCACCTTCTTTTGGCTTC<br>CAGATCCTCCTCAGAAATCAGC | AAGATGTGGAGAGTACGGGGTAG |
| mTmG | CTCTGCTGCCTCCTGGCTTCT | CGAGGCGGATCACAAGCAATA<br>TCAATGGGCGGGGGTCGTT |
| *Tek*-Cre | GCGGTCTGGCAGTAAAAACTATC | GTGAAACAGCATTGCTGTCAC TT |
| Sexing: *Sry* | TGGGACTGGTGACAATTGTC | GAGTACAGGTGTGCAGCTCT |
| Sexing: *Il3* | GGGACTCCAAGCTTCAATCA | TGGAGGAGGAAGAAAAGCAA |
| Sexing: *Myog* | TTACGTCCATCGTGGACAGC | TGGGCTGGGTGTTAGTCTTA |

Supplementary Table 1. Primers used for genotyping and sexing mice in the study.

#### *Maternal physiology cohort*

At 9-12 weeks postnatum, C57BL6/J (WT) virgin female mice were weighed and excluded from the study if <17 g. Female mice were then randomly allocated to the non-mated (Virgin) group or mated with stud males in the crosses described in Figure 6C. Females were monitored once daily between 08:00 and 10:00 for the presence of a vaginal plug. From e0.5, the dams were weighed then singly housed, as were Virgin group controls. On e14.5 (or equivalent single housing period for the Virgin group) animals were weighed then food was withdrawn at 6.00 pm until sacrifice between 8-10 am the following day (14-16 hr fast). On e15.5, the female mice were killed by terminal anaesthesia using ~0.8 mg pentobarbitol (Dolethal; Vetoquinol) per gram body weight, injected intra-abdominally. Upon cessation of the twitch reflex, mice were exsanguinated by cardiac puncture. Maternal and conceptus tissues were weighed and collected for processing. The following exclusion criteria were applied to all mice: (1) if the female mouse did not show evidence of a vaginal plug after 7 days spent with a stud male; (2) if a pregnant mouse carried <5 or >12 live conceptuses at e15.5; (3) if upon dissection the female mouse was found to have a confounding anatomical abnormality; (4) if the female mouse died during the experiment.

### Human samples

Samples were collected, staged and karyotyped by the facility. The 10 wkGA (8pcw), 15wkGA (13pcw) and 18wkGA (16pcw) placentas shown in Figure 1B and Figure 5G are respectively female, male and female, with normal autosomal karyotype.

### Determination of placental hormone levels in the maternal circulation

Levels of PRL3B1 and PRL8A8 were determined by Enzyme-linked immunosorbent assays provided by FineTest (EM1597, EM0798, Wuhan Fine Biotech Co, Ltd).

### Transcriptomic profiling of sorted cells from diverse placental lineages

#### *Cell sorting*

Embryos and placentas were harvested at e13.5 from Meox2-Cre females time-mated with *Dlk1*$^{-/+}$ (Mat); *Rosa26*$^{mTmG/\,mTmG}$ sires. *Meox2*-Cre embryos and placentas, distinguished by GFP expression, were visually identified using a Leica MZ10 stereomicroscope. Embryonic biopsies were collected for genotyping. Placentas were carefully sliced using sterile scalpels and subjected to a 1-hour 45-minutes incubation at 37 °C in a solution consisting of 0.1% Collagenase P (#11213865001, Roche-LifeScience,), 0.1% BSA (A9647, Merck) in PBS, to which DNase I (#07900, STEMCELL Technologies UK) was added. During the incubation period, samples were mechanically dissociated every 15-30 minutes, initially using a Pasteur pipette, and subsequently by passing the solution through needles of decreasing gauge sizes (18G, 21G and 23G). Cell suspensions were then filtered through 70-µm cell strainers attached to 50 ml conical tubes, then centrifuged at 1200 *g* for 5 minutes at 4 °C. Following this step, samples were washed with PBS supplemented with 0.1% BSA and treated with 1X RBC lysis buffer (#420301, BioLegend) in ddH2O for 5 minutes on ice with intermittent agitation for red blood cell removal. After an additional wash with PBS, the resulting cell pellets were resuspended in 400 µL of Cell staining buffer (# 420201, BioLegend) containing 4′,6-diamidino-2-phenylindole (DAPI, 1:5000, D9542, Merck). *Meox2*$^{+/+}$ placentas, which only express the membrane-localized tdTomato (mT) protein (TM+ samples), were used as negative controls. For the initial flow parameter setting experiment, wild-type (WT) stage-matched placental cells ($TM^{neg}$; $GFP^{neg}$) were used as additional negative controls. Cell sorting was conducted using a FACSAria™ III instrument, using the FACSDiva 8.0.1 (BD) software. The gating strategy was designed to exclude debris, doublets, and non-viable cells. The PE- and FITC-channels were used to specifically detect $TM^{+}$ and $GFP^{+}$ cells, respectively. For each sample, three distinct cell populations were selected: $TM^{+}$ ($PE^{high}$, $FITC^{neg}$); $GFP^{low}$ ($FITC^{low}$; $PE^{neg}$) and $GFP^{high}$ ($FITC^{high}$; $PE^{neg}$). Sorted cells were collected directly into TRIzol LS reagent (ThermoFisher) and stored at -80 °C for subsequent RNA extraction.

#### *RNA extraction*

Sorted cell samples were thawed at room temperature (RT), gently mixed by inversion and incubated at RT for 5 minutes. After adding chloroform in a ratio 3.75:1 TRIzol:Chloroform, samples were shaken vigorously, incubated for 5 minutes at RT and then centrifuged at 12000 *g* for 45 minutes at 4 °C. The upper aqueous phase containing the RNA was transferred to a new, clean tube. Samples were treated with RNase-free DNase I (NEB) at 37 °C for 10 minutes. Following the DNase treatment, EDTA was added to a final concentration of 5 mM before heat-inactivating the DNase I at 75°C for 10 minutes. RNA precipitation was achieved by adding 10 µg RNAse-free glycogen (AM9510, Invitrogen) and 100% isopropanol to the aqueous phase and allowing the RNA to precipitate overnight at -20 °C. On the following day, samples were centrifuged at 12000 *g* for 45 minutes at 4 °C to pellet the RNA. The resulting pellets were washed twice with 70% ethanol and centrifuged at 12000 *g* for 30 minutes at 4 °C, air dried and resuspended in RNase-free water. The purity of the extracted RNA was assessed using a Nanodrop spectrophotometer. Quantification of the RNA samples was performed using the RNA Quantification, high sensitivity assay kit for the QuBit (Q32852, Invitrogen). Sample integrity was evaluated using the Bioanalyser RNA 6000 Pico kit (5067-1513 , Agilent), following manufacturer's instructions, with the 2100 Expert Software.

#### *Library preparation and RNA sequencing*

Library preparation and sequencing were outsourced to Novagene. A total of 32 RNA samples with RNA Integrity Number (RIN) ≥6.8 (RIN Range: 6.8-9.2) were selected for library preparation and sequencing. These samples were derived from sorted placental cell populations, with 17 and 15 samples derived from $TM^{+}$ and the $GFP^{high}$ populations, respectively. For the $TM^{+}$ group, 9 placentas from male embryos (5 *Dlk1*$^{+/+}$; 4 *Dlk1*$^{+/-}$) and 8 from female embryos (4 *Dlk1*$^{+/+}$; 4 *Dlk1*$^{+/-}$) were chosen. For the $GFP^{high}$ group, the number of placentas from male and female embryos were 7 (3 *Dlk1*$^{+/+}$; 4 *Dlk1*$^{+/-}$) and 8 (4 *Dlk1*$^{+/+}$; 4 *Dlk1*$^{+/-}$), respectively. Whenever possible, samples were litter- paired and TM+ and GFP+ samples from the same placenta selected for analysis. Samples were derived from seven different litters. The cell count was 400000 cells per each of the $TM^{+}$ samples. For the the $GFP^{high}$ samples, the cell counts varied 31000 and 200000. The amount of total RNA sent for sequencing ranged from 54 and 300 ng per sample. RNA samples were converted into libraries using the Eukaryotic Transcriptome Library and subsequently sequenced using the paired-end 150 bp (PE150) platform provided by Illumina.

### Bioinformatic analyses

#### *RNA-seq data preprocessing*

Raw sequencing reads were trimmed and quality filtered using Trimmomatic (v. 0.39(11)) (parameters: LEADING:3; TRAILING:3; SLIDINGWINDOW:4:15; MINLEN:36; AVGQUAL:10), and then their quality was checked using FastQC (v. 0.11.9)[ https://www.bioinformatics.babraham.ac.uk/projects/fastqc/]. Reads were then mapped to the mouse reference genome (mm10) using the STAR aligner (v. 2.7.3a (12)) with default parameters. Read counts were estimated at gene level downloaded from Ensembl version 96) using the --quantMode GeneCounts option of STAR.

#### *Differential gene expression analysis*

Gene counts were normalized using the DESeq2 R/Bioconductor package (v. 1.28.1 (13)) having into account all factors (e.g. sex, litter, genotype) that may affect expression of each gene, followed by a regularized log transformation. In pairwise comparisons, Wald test P-values were adjusted by the Benjamini-Hochberg procedure for controlling the false discovery rate in multiple comparisons. To determine which genes were differentially expressed, we applied a filter requiring a gene to have mean normalized read count > 5 across all samples and either a 2-fold change in expression between GFP samples and TM samples, a 1.5-fold change in expression between Pat samples and WT samples, or a 1.5-fold change in expression between male (M) samples and female (F) samples, with a P-value < 0.05.

#### *Functional enrichment analysis*

Gene ontology (GO) enrichment analyses were performed using the clusterProfiler package (v. 3.16.1 (14)). P-values were calculated by hypergeometric distribution and adjusted for multiple hypotheses testing by the Benjamini-Hochberg procedure. Significantly enriched GO terms were selected according to an adjusted P-value < 0.01, and redundant GO terms were then removed by a semantic similarity-based approach implemented in the rrvgo package [https://ssayols.github.io/rrvgo]. Cell marker overrepresentation analysis was performed on genes in the upper quartile of expression in either GFP or Thermo TM samples using the enricher function of clusterProfiler with cell type markers retrieved from the PanglaoDB database (3). Significantly enriched cell types were selected according to an adjusted P-value < 0.01.

### Histology

Mouse placentas were fixed with 4% w/v paraformaldehyde (PFA, P6148, Merck) in Phosphate-Buffered Saline (PBS, BR0014G, Oxoid, Thermo Scientific) or Neutral Buffered Formalin (Merck, HT501128) overnight at 4°C and dehydrated through an increasing ethanol series the following day. Samples were stored at 4°C in 70% ethanol or dehydrated to 100% ethanol the day before the paraffin embedding. On the day of embedding, samples were incubated at room temperature (RT) with Histoclear II (National Diagnostics, HS202) (2 x 20 minutes for e9.5-e11.5; 2 x 35 minutes for e13.5) or Xylene (VWR) (2 x 45 minutes for e15.5-e18.5). This was followed by 3 x 1-hour incubations at 65°C with Histosec® (1.15161.2504, VWR). 5-7µm histological sections were cut using a Thermo HM325 microtome, mounted on Menzel-Gläser Superfrost®Plus slides (Thermo Scientific, J1810AMNZ) and used for Haematoxylin and Eosin (H&E) staining, *in situ* hybridisation (ISH), RNAscope, immunohistochemistry (IHC) and immunofluorescence (IF).

#### *Immunohistochemistry and immunofluorescence*

IHC on histological sections was performed as previously described (15). For IHC, unmasking was achieved by boiling the histological sections with 10 mM tri-sodium citrate buffer pH 6 for 20 minutes. Detection of the proteins was achieved by incubating the histological sections overnight at 4°C with the primary antibodies described in Supplementary Table 2, and detected with biotinylated goat α-rabbit, goat α-mouse or horse α-goat secondaries (BA-1000, BA-9200 and BA9500, Vector Laboratories, 1:300), followed by 1-hour incubation at room temperature with Vectastain® Elite ABC-HRP kit (PK-6100, Vector Laboratories). Staining was achieved through colorimetric reaction using DAB Peroxidase Substrate Kit (SK-4100, Vector Laboratories). Slides were lightly counterstained with Mayer's Haematoxylin (MHS16, Merck) or Nuclear Fast Red (H-3403-500, Vector Laboratories) and mounted using DPX (Merck, 06522).

For IF, unmasking was performed as above except with Tris-EDTA buffer pH 9 [10 mM Tris Base, 1 mM EDTA, 0.05% Tween 20]. Histological sections were incubated for 1hr in blocking buffer [5% Blocker Casein (Thermo #37528) with 0.05% Tween 20 in PBS] then incubated overnight at 4°C with the primary antibody (Supplementary Table 2) in blocking buffer. Sections were incubated with the appropriate fluorescence conjugated secondary antibody [horse α-rabbit DyLight® 594 (DI-1094 Vector Laboratories), donkey α-goat Alexa-fluor-488 or 647, (A11055, A21447, Invitrogen), donkey α-rat Alexa-fluor-488 (A48269, Invitrogen), goat α-mouse DyLight® 488 (ab96871, Abcam), goat α-mouse Alexa-fluor-594 (A11032, Invitrogen), all 1:300] for 1 hour. Autofluorescence was quenched using the Ready Probes Tissue Autofluorescence Quenching kit (R37630, Invitrogen), then mounted using VECTASHIELD® Antifade Mounting Medium with DAPI (H-1200-10, Vector Laboratories). Isotype controls for each antibody showed no staining under identical conditions.

| Target | Product | Working concentration |
|---|---|---|
| Mouse DLK1 | Abcam ab210471 | 1:250 |
| Human DLK1 | R&D Systems #AF1144 | 1:100 |
| Mouse EMCN | SantaCruz sc-65495 | 1:200 |
| Mouse SMA (1A4) | Invitrogen MA1-06110 | 1:4000 |
| Human CD31 | Abcam ab28364 | 1:50 |
| Human CK8 clone TROMA-1 | Merck-Millipore MAB7329 | 1:250 |
| Mouse CK7 | Abcam ab181598 | 1:250 |
| Mouse FOXO1 | Cell Signalling Technologies #2880 | 1:100 |

Supplementary Table 2. Primary antibodies used in this study.

#### *Imaging*

Images of histological sections were acquired using a NanoZoomer HT (Hammamatsu). Fluorescence images and higher magnification bright-field images were acquired using a Zeiss Axioplan II microscope with a Luminera 3 digital camera and INFINITY ANALYSE software v6.5.6 (Luminera), or Zeiss AXIO Observer Z1 with a Teledyne-Photometrics Iris 9 camera, captured with Micromanager v2 software (16). Images were combined into panels using *Adobe Photoshop 26.4.0 release 2024 (Adobe)*.

#### *Stereological estimation of pituitary volumes and cell proportions.*

Samples were blinded prior to stereological investigations. Placental volume was estimated using the Cavallieri method (17). Briefly, the placenta was exhaustively sectioned sagitally then 7μm sections were collected at regular, non-overlapping intervals throughout the organ from a random starting point and stained with H&E. Images of each H&E-stained section were acquired using a NanoZoomer HT

(Hamamatsu) and processed using NDP.view2 software (Hamamatsu) to calculate the total cross sectional area (CSA) 20 times per placenta, which was then converted to volume.

Cell proportions were estimated as described previously (18). Briefly, for each animal 6 non-overlapping H&E sections from the lateral and midline of the placenta at 200x magnification were overlaid with a point counting grid. The cellular identity of the cell beneath the points were identified as maternal decidua, trophoblast giant cells (T-GC), glycogen cells (GlyT), spongiotrophoblast (SpT) and chorionic plate (CP). Cell types of the labyrinth could not be distinguished in H&E staining, so labyrinthine trophoblast, fetal endothelial cells and mesenchymal cells of the labyrinth were combined into a single category (LAB). ~2000-8000 cells were classified per placenta depending on the developmental stage.

Fetal endothelial CSA was determined from sections stained by IHC for the endothelial marker endomucin (EMCN). Images of a randomly chosen midline section from each placenta were acquired using a NanoZoomer HT (Hamamatsu) and processed using NDP.view2 software (Hamamatsu). The freehand annotation tool was used to draw around the periphery of endothelial cells, as generated by the EMCN staining, generating a CSA of each vessel. Between 150-200 vessel CSAs were collected per section and the mean and median CSA calculated per placenta.

Quantification of FOXO1-positive endothelial cells was performed using QuPath (19). Specifically, two images of each DAPI+/EMCN+(GFP)/FOXO1+(RFP) stained placenta were captured at 20x magnification (one proximal to the junctional zone and one at the chorionic plate). Tri-colour images were uploaded onto QuPath without further processing. The labyrinth in each image was selected by reference to the EMCN+ region, then all nuclei counted in this region using the cell detection tool at fixed threshold. Classifiers were used to define the FOXO1+ nuclei (default parameters for nuclei in the RFP channel, set threshold) and EMCN+ cells (default parameters for cytoplasm in the GFP channel, set threshold). % counts were the average of the values obtained from the two images, as a proportion of cells defined by DAPI+ nuclei.

### Placenta lysate preparation and Western blotting

Tissue lysates were prepared from whole placentas collected at e13.5. Placentas were snap frozen in liquid nitrogen and stored at -70°C until further use. To obtain tissue lysates, each frozen placenta was placed in a pre-cooled Douce homogeniser containing 700μl of TBS (0.4 M NaCl, 0.1 M Tris pH7.5) and protein loading buffer (EC-887, National Diagnostics) diluted at 1:1 ratio with the total volume. 1% β-mercaptoethanol was added to the protein loading buffer immediately prior use. Placentas were homogenised by applying 10 strokes with the loose pestle and 10 strokes with the tight pestle. Samples were placed in individual 1.5ml Bioruptor tubes (C30010016. Diagenode) and sonicated at 4°C for 30 seconds 25 times (with a 30 second interval between cycles). Tissue lysates were then triturated by passing five times through a disposable 26-gauge needle (AN2623R1, Terumo) attached to a 1 ml syringe. Sonication and trituration was repeated. 16 μl of each protein lysate was denatured at 95°C for 10 minutes and subjected to SDS-PAGE. Proteins were transferred to a nitrocellulose membrane (PB3310, Invitrogen) and blocked for 1h at room temperature in either 3% BSA (bovine serum albumin) or 3% dried milk in TBS-T (TBS with 0.1% Tween 20). Membranes were then incubated overnight at 4°C with individual primary antibodies: Rabbit anti-FOXO1 (2880, Cell Signaling Technology, 1:500 in 3% BSA in TBS-T) and rabbit anti-CD31 (GTX130274, GeneTex, 1:800 in 3% dried milk in PBS-T). Membranes were incubated for 1h at room temperature with a secondary antibody (anti-rabbit HRP, P0448 Agilent). Protein detection was performed by chemiluminescence using Pierce ECL Plus Substrate (32132, Thermo Scientific™) according to manufacturer's instructions. Anti-alpha tubulin (T5168, Merck,1:10000) levels were used to normalise the total level of proteins. Membranes were imaged using a ChemiDoc™ Imaging System (BIO-RAD), and protein quantification was performed using Image Lab 6.1 software (BIO-RAD).
